# Heterologous expression of *Dehalobacter* spp. respiratory reductive dehalogenases in *Escherichia coli*

**DOI:** 10.1101/2021.10.06.463429

**Authors:** Katherine J. Picott, Robert Flick, Elizabeth A. Edwards

## Abstract

Reductive dehalogenases (RDases) are a family of redox enzymes that are required for anaerobic organohalide respiration, a microbial process that is useful in bioremediation. Structural and mechanistic studies of these enzymes have been greatly impeded due to challenges in RDase heterologous expression, primarily because of their cobamide-dependence. There have been a few successful attempts at RDase production in unconventional heterologous hosts, but a robust method has yet to be developed. In this work we outline a novel respiratory RDase expression system using *Escherichia coli* as the host. The overexpression of *E. coli*’s cobamide transport system, *btu*, and RDase expression under anaerobic conditions were established to be essential for the expression of active RDases from *Dehalobacter* - an obligate organohalide respiring bacterium. The expression system was validated on six RDase enzymes with amino acid sequence identities ranging from >30-95%. Dehalogenation activity was verified for each RDase by assaying cell-free extracts of small-scale expression cultures on various chlorinated substrates including chloroalkanes, chloroethenes, and hexachlorocyclohexanes. Two RDases, TmrA from *Dehalobacter* sp. UNSWDHB and HchA from *Dehalobacter* sp. HCH1, were purified by nickel affinity chromatography. Incorporation of both the cobamide and iron-sulfur cluster cofactors was verified, and the specific activity of TmrA was found to be consistent with that of the native enzyme. The heterologous expression of respiratory RDases, particularly from obligate organohalide respiring bacteria, has been extremely challenging and unreliable. Here we present a relatively straightforward *E. coli* expression system that has performed well for a variety of *Dehalobacter* spp. RDases.

**IMPORTANCE:** Understanding microbial reductive dehalogenation is important to refine the global halogen cycle and to improve bioremediation of halogenated contaminants; however, studies of the family of enzymes responsible are limited. Characterization of reductive dehalogenase enzymes has largely eluded researchers due to the lack of a reliable and high-yielding production method. We are presenting an approach to express reductive dehalogenase enzymes from *Dehalobacter*, a key group of organisms used in bioremediation, in *E. coli*. This expression system will propel the study of reductive dehalogenases by facilitating their production and isolation, allowing researchers to pursue more in-depth questions about the activity and structure of these enzymes. This platform will also provide a starting point to improve the expression of reductive dehalogenases from many other organisms.

## INTRODUCTION

A wide array of naturally produced halogenated organic compounds, or organohalides, are present in the environment. Organohalides also make up some of the most pervasive and concerning anthropogenic soil, sediment, and groundwater contaminants. Anaerobic organohalide-respiring bacteria (OHRB) can readily degrade a number of these pollutants, such as chlorinated ethenes and ethanes, by using them as terminal electron acceptors in their cellular respiration (1, 2). The respiration process removes halogen atoms from organohalide substrates via reductive dehalogenation, changing the properties of the parent substrate, often detoxifying or transforming them to more degradable substrates for other organisms. OHRB were first described in 1990 (1), and quite rapidly became a staple tool in commercial bioremediation. In particular, obligate OHRB *Dehalococcoides* and *Dehalobacter* are highly abundant in bioremediation cultures targeting chlorinated contaminants (3, 4). Obligate OHRB have very restricted metabolism and can only grow on organohalide substrates while facultative OHRB are flexible and can utilize a variety of electron acceptors (2). All OHRB use reductive dehalogenase enzymes (RDase) discovered in 1995 as part of their electron transport chain to perform these reduction reactions (5). RDases are key targets for studying the mechanism of OHRB respiration and remediation, and on their own RDases offer an intriguing tool for industrial applications.

RDases form a large family of oxygen-sensitive enzymes that rely on two iron-sulfur clusters ([4Fe-4S]) and a cobamide, commonly cobalamin or vitamin B_12_, as cofactors to perform catalysis; RDases are well described in previous reviews (2, 6). There are two broad categories: respiratory RDases from OHRB, and catabolic RDases which are catalytically similar but perform a different cellular function (7, 8). Here we will focus on respiratory RDases as they represent an unusual metabolism and are more relevant to bioremediation. All RDases remove halogen atoms from their substrate through either a hydrogenolysis reaction, where one halogen is substituted with a hydrogen, or a β-elimination reaction, where two halogen atoms across a single bond are removed (this may also occur with a halogen and a hydrogen in a non-energy yielding process). Respiratory RDases also hold a twin arginine translocation (TAT) signal peptide to localize them to the periplasmic membrane, though these are cleaved off in the mature protein. While the RDase family has wide sequence variation, many of the described RDases are from *Dehalococcoides* and *Dehalobacter* and use chloroethenes, chloroethanes, or chlorobenzenes as substrates. Furthermore, obligate OHRB encode dozens of unique *rdhA* genes in their genomes (9); although, majority are not expressed in culture and have unknown functions. Approximately 25 unique RDases have had their substrates experimentally determined (see Table S1 for details), yet there are almost 1000 unstudied *rdhA* (RDase homologous genes) sequences in the Reductive Dehalogenase Database (9), and many more in metagenomic surveys such as the ones described in (10, 11). The number of identified *rdhA* sequences continues to climb with more sequencing, yet functional characterization methods lag far behind.

The majority of the functionally characterized RDases were purified from their endogenous (or native) producers (Table S1) through extensive methods (5, 12, 21–23, 13–20). Obtaining enough biomass for such purifications is not a trivial undertaking and the purification methods under anaerobic conditions are tedious. Because most OHRB are difficult to grow as they require toxic, poorly soluble, and often volatile halogenated substrates, small scale partial purification using blue native-polyacrylamide gel electrophoresis (BN-PAGE) coupled with activity assays and proteomic analysis has proven very useful for ascribing substrates to specific RDases (24–29). All of the above characterization studies are confined both to lab-cultivatable organisms and to actively expressed genes. Table S1/SI Text 1 provides a list of currently known RDases.

To address the limitations of purifying RDases from native producers, many research groups have attempted heterologous expression, but the complex cofactor requirements, which are crucial for RDase activity, have hindered efforts. We compiled a list of all expression studies (Table S2/SI Text 1). The first published expression attempt was in 1998, where PceA from *Sulfurospirillum multivorans* was expressed in *Escherichia coli* but no activity detected (30). Subsequent trials in *E. coli* involved denaturation and cofactor reconstitution (31, 32). It was not until 2014 when the first activity from heterologously expressed RDases were from the cobamide producing host *Shimwellia blattae*; however, these enzymes were unable to be purified using a Strep affinity tag (33). Finally, twenty years after their discovery, the first RDase crystal structures were solved: one of PceA from *S. multivorans*; obtained from the native producer with a mutation to incorporate an affinity tag (34), and the catabolic RDase NpRdhA from *Nitratireductor pacificus* pht-3B; expressed in the cobalamin producing *Bacillus megaterium* (8). The two crystal structures confirmed that the cobamide cofactor is deeply buried within the enzyme and makes numerous interactions with the peptide sidechains suggesting that it may serve a structural role for proper enzyme folding (8, 34). Most typical heterologous hosts, such as *E. coli*, neither produce cobamides *de novo* nor import the molecule under basal conditions. Since the crystal structures, there have been a couple successful reports of catalytically active RDase expression all using hosts *S. blattae* and *B. megaterium*, both with full cobamide biosynthetic pathways (35, 36). However, less conventional hosts tend to have lower protein yields and have less standardized protocols. The preferable host would be *E. coli* as it grows quickly, is highly engineered to obtain high protein yield, and is amenable to modern molecular biology tools. Thus far, expression attempts using *E. coli* have required enzyme denaturation and refolding for cofactor reconstitution to obtain activity (32, 37, 38), but these methods are too unpredictable and unreproducible to be used as a standard expression system. Although developing a generalizable RDase expression system has been a goal in the field for decades, to date, no respiratory RDase has been directly produced from *E. coli* in its active state.

Although *E. coli* does not synthesize cobalamin *de novo*, it, like many other organisms, possesses the vitamin B_12_ salvage pathway (*btu*) to take up cobalamin from the environment (39). This salvage pathway consists of a series of transport proteins: BtuB, a TonB-dependent transporter protein on the outer membrane to import cobalamin into the periplasm; BtuF, a periplasmic protein that will bind free cobalamin; BtuCD, an ABC-transporter that associates with BtuF to import cobalamin into the cytoplasm, a cartoon representation can be found in Figure 1B (40–46). Expression of the *btu* operon is poor under typical growth conditions but can be enhanced with the use of ethanolamine-based media, which forces *E. coli* to use a cobalamin-dependent enzyme, or by induced expression (44, 47). Recently, the Brooker lab developed an expression system for cobalamin-dependent *S*-adenosylmethionine methylases that expressed the entire *btu* operon from *E. coli* under arabinose induction in the vector *pBAD42-BtuCEDFB* (44). The application of this vector greatly enhanced the solubility and activity of several methylases tested (44). Similarly, the catabolic RDase, NpRdhA, had an slight increase in yield and cofactor incorporation when co-expressed with just BtuB (48). Given the success of these expression systems and the hypothesis that the cobamide cofactor supports proper RDase folding, we predicted that implementation of *btu* expression could result in generation of active RDases in *E. coli*.

**Figure 1.**
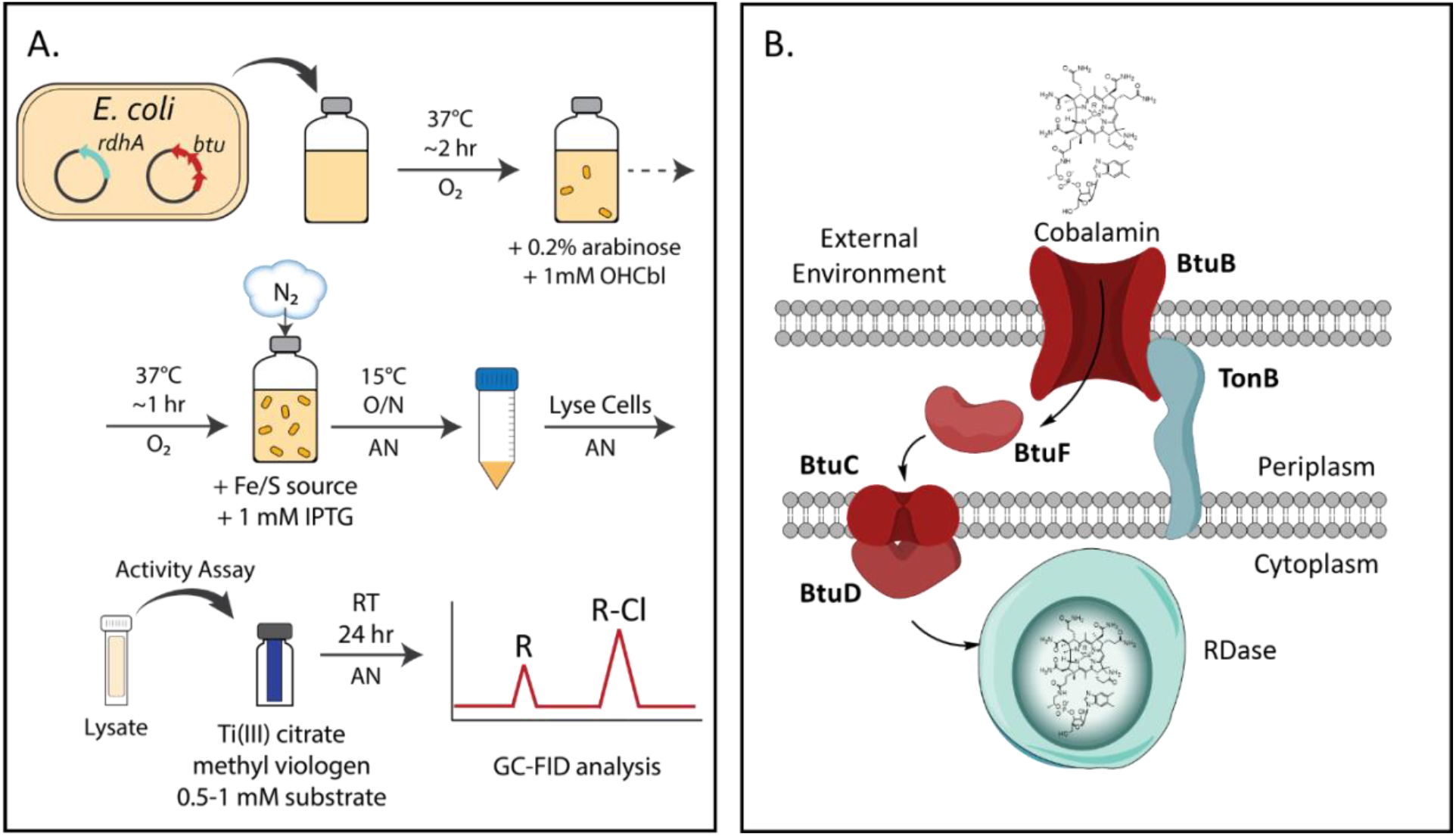
(A) Schematic of the RDase expression system and small-scale expression tests to determine production of the active RDases. (B) Cartoon of the Btu pathway used for cobalamin incorporation in RDase expression, figure adapted from (44). O_2_ = aerobic conditions, AN = anaerobic conditions, O/N = overnight, GC-FID = gas chromatography with flame ionization detection.

In this work, we employ the *pBAD42-BtuCEDFB*, provided by the Booker lab, in several strains of *E. coli* with enhanced iron-sulfur cluster production and maintenance, for the heterologous expression of several respiratory RDases from *Dehalobacter* spp. This expression system is a tremendous steppingstone to advance knowledge about the RDase enzyme family and organohalide respiration, and to facilitate future development in bioremediation.

## Materials and Methods

All chemicals and primers were purchased from Sigma-Aldrich (St. Louis, MO, USA) except for antibiotics, terrific broth (TB) media, and Bradford reagent which were purchased from BioRad (Hercules, CA, USA). All gases were supplied from Linde Canada Inc. (Mississauga, ON, Canada). Synthetic genes and the *pET21-hchA* plasmid were synthesized by Twist Bioscience (San Francisco, CA, USA) using codon optimized sequences for TmrA (WP_034377773), CfrA (AFV05253), DcrA (AFV02209), and HchA (IMG Accession: 2823894057). DNA isolations and purifications were done with the QIAprep Spin Miniprep Kit and the QIAquick PCR Purification Kit, and nickel-nitrilotriacetic acid (Ni-NTA) agarose resin was purchased from Qiagen (Hilden, Germany). PCR reagents, restriction enzymes, Gibson Assembly master mix, the 1 kb DNA ladder, and *Escherichia coli* DH5α were purchased from New England Biolabs (Ipswich, MA, USA). Sanger sequencing was performed by The Centre for Applied Genomics at SickKids Hospital (Toronto, ON, Canada). The *pBAD42-BtuCEDFB* plasmid was generously provided by the Booker Lab (Pennsylvania State University, PA, USA). *E. coli* BL21(DE3) Δ*iscR* was obtained from Patrick Hallenbeck (49, 50), *E. coli* BL21(DE3) *cat-araC-*P_BAD-*suf*_, Δ*iscR::kan*, Δ*himA::*Tet^R^, abbreviated to *E. coli ara-Suf* Δ*iscR*, and *E. coli* SufFeScient were generously provided by the Antony and Kiley Labs (St. Louis University School of Medicine, MO, USA; University of Wisconsin-Madison, WI, USA) (51). The empty *p15TV-L* plasmid is available at Addgene #26093. All buffers used in protein extractions and purifications were purged with N_2_ gas prior to use.

### Plasmid Construction and Transformation

Full *tmrA*, *cfrA*, and *dcrA* genes were codon-optimized and ordered as synthetic gene blocks (Supplemental Text 1). Two primer sets were designed for amplification of the genes with extensions to introduce a 15 bp overlap sequence for Gibson Assembly into the linearized *p15TV-L* plasmid. One primer pair amplified the TAT signal peptide sequence and introduced a C-terminal x6His tag extension in case the N-terminal tag was cleaved with the endogenous TAT processing machinery, this pair was only designed for TmrA amplification. The second primer pair amplified the gene without the TAT sequence to truncate the cloned gene, the gene began at the previously observed TAT cleavage site (14). Primer sequences and their descriptions are in Table S3/SI Text 2. Genes were amplified by PCR using the synthetic genes as the templates. The *p15TV-L* plasmid was linearized by BseRI digestion. The linear plasmid and the amplified gene products were purified with a QIAquick PCR Purification kit. The amplified gene product and the linearized plasmid were ligated by Gibson Assembly, transformed into *E. coli* DH5α competent cells via chemical transformation, and plated on carbenicillin (100 ng/mL) selection plates.

Positive transformants were picked, grown up, and their plasmids were extracted. The plasmids were screened by PCR amplification of the desired gene and Sanger sequencing using universal T7 primers. The sequencing-confirmed plasmids were transformed into both *E. coli* Δ*iscR* and *E. coli ara-Suf* Δ*iscR* strains with and without *pBAD42-BtuCEDFB*; those with *pBAD42-BtuCEDFB* were selected with carbenicillin and spectinomycin (50 ng/mL) plates. The plasmid *p15TVL-tmrA* was also transformed into *E. coli* BL21(DE3) Lobstr and *E. coli* SufFeScient. The codon-optimized *hchA* gene without the TAT signal sequence, as predicted by SignalP-5.0 (52), was ordered in *pET-21* with a C-terminal x6His tag. The genes for DHB14 and DHB15 had been previously amplified from genomic DNA isolated from the ACT-3 mixed culture and cloned into *p15TV-L* with their TAT signal sequences by the Yakunin and Savchenko Labs (University of Toronto, Toronto, ON, Canada). The plasmids *pET21-hchA, p15TVL-DHB14,* and *p15TVL-DHB15* were transformed into *E. coli* Δ*iscR* with *pBAD42-BtuCEDFB*. Descriptions of all expression plasmids are included in Table S4/SI Text S2, and descriptions of the *E. coli* strains used are in Table S5/SI Text S2.

### Small-Scale Protein Expression and Lysis

Full schematic of small-scale expression and lysate activity assays is shown in Figure 1. An aliquot (200 µL) of starter cultures containing the desired expression plasmids were used to inoculate 20 mL of TB media with the appropriate antibiotics in 160 mL serum bottles, or 250 mL Erlenmeyer flasks if full expression was aerobic. In initial tests to establish the expression system, cultures in the serum bottles were grown anaerobically by sealing the bottle and purging the media with N_2_ prior to inoculation, these were then treated the same as the aerobic cultures (same induction and incubation times) though the cell density tended to be lower in these samples. After further optimization the finalized expression system is described here. Bottles and flasks were kept aerobic for growth, and the cultures were incubated at 37°C, 180 rpm until the optical density at 600 nm (O.D._600_) reached 0.4-0.6. Cultures were supplemented with a final concentration of 1 µM hydroxycobalamin and *btu* expression was induced with a final concentration of 0.2% (w/v) arabinose (regardless of whether *pBAD42-BtuCEDFB* was present in the culture strain). Cultures were incubated again at 37°C, 180 rpm until the O.D._600_ reached 0.8-1. All cultures were put on ice, cultures in serum bottles were stoppered, crimped, and the headspace was purged with N_2_ gas for 10 min to remove excess O_2_, cultures in flasks remained aerobic. All cultures were supplemented with final concentrations of either 50 μM ammonium ferric sulphate used during TmrA expression optimization, or 50 μM cysteine and 50 μM ammonium ferric citrate used with all subsequent cultures to align with recommended iron-sulfur sources though no change in activity was observed (53), and RDase expression was induced using 1 mM isopropyl β-D-1-thiogalactopyranoside (IPTG). Cultures were incubated at 15°C, 180 rpm overnight. Expression of TmrA under different conditions tested is in Figure S1/SI Text 3.

After incubation, all cultures were handled in a Coy anerobic glovebox with a supply gas composition of 10% H_2_, 10% CO_2_, and 80% N_2_. Cultures were harvested by centrifugation at 6000 xg (Avanti-JE SN:JSE07L45, Beckmann Coulter) and 4°C for 10 min. Supernatants were discarded and the pellets were resuspended in 1 mL of Buffer A (50 mM Tris-HCl (pH 7.5), 150 mM NaCl, 5% glycerol, 0.1% Triton X-100) with 1 mM tris(2-carboxyethyl)phosphine (TCEP) and Protease Inhibitor Cocktail (PIC; final concentrations: 1 mM benzamidine, 0.5 mM phenylmethylsulfonyl fluoride) added fresh at time of use. In some cases, Triton X-100 in Buffer A was replaced with either 1% 3-((3-chloamidopropyl)dimethylammonio)-1-propanesulfonate (CHAPS) or 1% digitonin. The suspended pellet was transferred to a 2 mL o-ringed microcentrifuge tube with 50 mg of glass beads (150-500 μm). The cells were lysed via bead beating by vortexing the tubes for 2 min and resting on ice for 1 min, repeated 3 times. The lysate was clarified by centrifugation at 20 000 xg (Thermo Scientific microcentrifuge, serial: 1931100817155), 4°C for 5 min. The soluble lysate fraction was used for subsequent activity assays. The positive controls using ACT-3, a mixed culture enriched maintained on either chloroform or 1,1,1-trichloroethane as electron donor, were prepared the same way only 0.5 mL of buffer was used for resuspension.

### TmrA and HchA Large-Scale Protein Expression and Purification

An overnight start culture (5 mL) of either *E. coli ara-Suf* Δ*iscR p15TVL-tmrA* + *pBAD42-BtuCEDFB* or *E. coli* Δ*iscR pET21-hchA* + *pBAD42-BtuCEDFB* was used to inoculate 1 L of TB medium with appropriate antibiotics in a 2 L glass bottle (Fischer Scientific), that was kept aerobic for growth with a foil cap. The culture was incubated at 37°C, 150 rpm until the O.D._600_ reached 0.4-0.6, 3 μM hydroxycobalamin was supplemented to the culture and 0.2% (w/v) arabinose was used to induce *btu* expression. This was incubated at 37°C, 150 rpm again until the O.D._600_ reached 0.8-1, after which the culture was put on ice, sealed with a rubber septa and cap, and the headspace was purged with N_2_ gas for 1 hr to make anaerobic. The culture was supplemented with 50 μM cysteine and 50 μM ammonium ferric citrate, and RDase expression was induced with 1 mM IPTG. The culture was then incubated overnight at 15°C, 150 rpm.

The expression culture was handled in a Coy anaerobic glovebox or kept sealed in an airtight vessel for all of the following steps. The culture was harvested by centrifugation at 6000 xg, 4°C for 15 min (Avanti-JE SN:JSE07L45, Beckmann Coulter). The supernatant was discarded, and the pellet was resuspended in 5 mL of Binding Buffer (Buffer A, 30 mM imidazole) per 1 g of wet cell weight with 1 mM TCEP and PIC added fresh to the buffer from concentrated stocks. The cells were lysed with the addition of 10x Bug Buster concentrate (Millipore), at the proper dilution, 0.3 mg/mL lysozyme, 1.5 µg/mL DNase, and 10 mM MgCl_2_ (final concentrations). The lysate was incubated at RT shaking at 50 rpm for 20 min. The lysate was clarified by centrifugation at 29 000 xg, 4°C for 20 min (Avanti-JE SN:JSE07L45, Beckmann Coulter).

A 1 mL Ni-NTA resin in a glass gravity-flow column (Econo-Column^®^, BioRad, USA) was set up inside a Coy anaerobic glovebox with an atmosphere of 100% N_2_. Ni-NTA (1-2 mL) resin was purged with N_2_ prior to use. The column was washed with 5 column volumes (CVs) of ddH_2_O and conditioned with 5 CVs of Binding Buffer. The clarified cell lysate was loaded onto the column, flow through was collected for subsequent analysis. The column was washed with 20 CVs of Binding Buffer until no protein was detected in the eluent by Bradford reagent. The RDase was eluted from the column using Elution Buffer (Buffer A, 300 mM imidazole). Collected fractions underwent a buffer exchange into Buffer A (without imidazole) with 1 mM TCEP, to remove the imidazole and the protein was concentrated using a 30 kDa cut-off Millipore filter tube. The concentrated protein was quantified using a Bradford assay and distributed into aliquots for flash freezing and storage in liquid N_2_. All steps of the purification were imaged and analysed by SDS-PAGE, the purity of the final enriched enzyme was estimated by band density using Image Lab v6.1 Software (Bio-Rad Laboratories, Inc.); the SDS-PAGE of TmrA and lane density for purity analysis is shown in Figure S2 and protein yield is in Table S6 of SI Text 3.

### Dechlorinating Activity Assays

Initial protein activity assays to test Btu expression and effect of oxygen were carried out in 2 mL reaction volumes, all subsequent assays were done at a volume of 500 μL to conserve materials. All assays were carried out in glass vials with no headspace and sealed with polytetrafluoroethylene (PFTE)-lined caps, these were set up and incubated under anaerobic conditions in a Coy glovebox with a supply gas composition of 10% CO_2_, 10% H_2_, and 80% N_2_. The assay was carried out in a buffer of 50 mM Tris-HCl (pH 7.5), 2 mM Ti(III) citrate, and 2 mM methyl viologen. Enzyme was added in the form of either 50 μL soluble cell lysate (100 μL for 2 mL reaction volumes) or 0.3-2 μg purified TmrA/HchA. All substrates, except the hexachlorocyclohexane (HCH) isomers, were supplemented from saturated water stocks to final concentrations of either 0.05 mM (in the case of perchloroethene), 0.5 mM, or 1 mM using Hamilton glass syringes. HCH was added from a 10 mM stock solution in DMSO to a final concentration of 0.3 mM. Lysate assays were kept at RT overnight (18-24 hr), purified enzyme assays were kept at RT for 1 hr and then were stopped by acidification as described below.

All assays were stopped by transferring 0.4 mL (1 mL was taken for reactions with 2 mL total volumes) of the reaction volume into acidified water (pH < 2 using HCl) for a total sample volume of 6 mL; this was sealed in a 10 mL vial for headspace analysis by gas chromatography with flame ionization detection (GC-FID). The assays were all run using the same GC-FID separation method on an Agilent 7890A GC instrument equipped with an Agilent GS-Q plot column (30 m length, 0.53 mm diameter). Sample vials were equilibrated to 70°C for 40 min in an Agilent G1888 autosampler, then 3 mL of the sample headspace was injected (injector set to 200°C) onto the column by a packed inlet. The flow rate was 11 mL/min of helium as the carrier gas. The oven was held at 35°C for 1.5 min, raised at a rate of 15°C/min up to 100°C, the ramp rate was reduced to 5°C/min until the oven reached 185°C at which point it was held for 10 min. Finally, the oven was ramped up to 200°C at a rate of 20°C/min and held for 10 min. The detector was set to a temperature of 250°C. Data were analyzed using Agilent ChemStation Rev. B.04.02 SP1 software, and peak areas were converted to liquid concentration using standard curves (external calibration) for each compound. An example chromatogram is shown in Figure S3/SI Text 3.

Three types of negative controls for the enzyme assays were used: enzyme-free controls, deactivated enzyme controls, and free cobalamin (cobalamin only; no enzyme) controls. Enzymes were deactivated by boiling lysate or purified sample for 5-10 min, this was only done for one representative enzyme for each substrate to verify there was no abiotic reaction from the *E. coli* lysates. Reduction using free cobalamin was tested using a final assay concentration of 0.2 μM cobalamin for lysate assays, and at an equimolar concentration to the enzyme when used as a negative control in the purified enzyme assays. ACT-3 mixed culture was used as a positive control where possible, using 50 μL of soluble cell lysate, similar to the heterologously expressed enzymes. All enzyme lysates were tested with six replicates (two lysates from independently grown expression cultures with three assays each) on their known substrates to confirm activity, and a minimum of duplicates on additional substrates (two lysates from independently grown cultures). Negative controls for α-HCH were only done as single samples due to limited substrate. The purified TmrA/HchA preparations were tested in triplicates. The number of replicates is indicated where the data are presented, all of the individual sample raw data are shown in excel Tables S7-S11.

### Cofactor Quantification

Iron content of purified TmrA and HchA was quantified using a ferrozine method as previously described (49, 54, 55). A range of 0.5-20 nmol FeCl_3_ was used to create a standard curve. Protein samples were boiled for 5 mins; both the samples and standards were mixed with 300 µL iron-releasing agent (1.4 M HCl, 0.3 M KMnO_4_) and heated at 60°C for 30 min. This was brought to RT and then mixed with 150 µL iron-binding agent (6.5 mM ferrozine, 6.5 mM neocuproine-HCl, 2.5 M ammonium acetate, 1 M ascorbic acid). The absorbance was measured at 560 nm and iron content in the protein sample was determined by the standard curve and adjusted with the purity of TmrA/HchA to get an occupancy ratio that was not skewed by the impurities.

Cobalamin was quantified using two separate methods, in both cases cyanocobalamin at a concentration range of 0.1-10 µM (final concentration) was used for a standard curve. The first method was used as previously described with some modifications (35, 44, 56). The standard samples and protein samples (50 μL) were mixed with 15 mM KCN to a volume of 750 µL and were heated at 80°C for 10 min. The samples were clarified by centrifugation and their absorbance was measured at 367 nm and at 650 nm to normalize to the baseline. The second method was previously described (57, 58); briefly, samples were mixed with 10 mM KCN in 80% (v/v) methanol and acidified using 3% (v/v) acetic acid. Samples were then heated at 80°C for 1 hr with slow shaking. Samples were clarified by centrifugation, the supernatant was transferred and dried, and the extract was resuspended in ddH_2_O such that the concentration was unaltered. These samples were analysed using ultra-high performance liquid chromatography-mass spectrometry (UHPLC-MS). For both methods the cobalamin occupancy was adjusted for the purity of the enzymes.

The samples analyzed by UHPLC-MS were separated on a Thermo Scientific Ultimate 3000 UHPLC instrument equipped with a Thermo Scientific Hypersil Gold C18 column (50 × 2.1 mm, 1.9 µm particle size) and a guard column held at 30°C. The sample (10 µL) was injected by an autosampler held at 5°C. Eluent A was 5 mM ammonium acetate in water, eluent B was 5 mM ammonium acetate in methanol, the flow rate was set to 300 µL/min. The separation method used started with a mix of 2% eluent B for 0.46 min, then held a linear gradient to 15% B from 0.46-0.81 min; a linear gradient to 50% B from 0.81-3.32 min; a linear gradient to 95% B from 3.32-5.56 min; 95% B was held until 7.5 min; finally, the composition was reduced to 2% B with a linear gradient from 7.5-8 min, and held at 2% B from 8-12 min. The eluent was introduced to a Thermo Scientific Q Exactive mass spectrometer with a heated electrospray ionization (HESI-II) probe at a spray voltage of 3.5 kV, and capillary temperature of 320°C. Data collection was done in positive ion modes with a scan range of 1000-2000 m/z, mass resolution of 70 000, an automatic gain control target of 3×10^6^, and a maximum injection time of 200 ms. Data were processed using Thermo Xcalibur Qual Browser v3.1 software, cyanocobalamin was detected at an [M + H] ^+^ of 1355.5747 (ppm error = 5), the peak area was used for quantification.

## RESULTS & DISCUSSION

### Developing the expression system in *E. coli*

The expression system was developed using TmrA, a chloroalkane-reducing RDase from *Dehalobacter* sp. UNSWDHB. It was chosen because it was previously expressed in its active form in *B. megaterium* (35). TmrA was cloned without its TAT signal peptide into the *p15TV-L* expression vector. This construct expressed TmrA with an N-terminal x6His tag under IPTG control. The expression vector was introduced into *E. coli* Δ*iscR*, a strain engineered for enhanced [4Fe-4S] production by deleting the *isc* (iron-sulfur cluster) operon repressor (49, 50). Small-scale expression cultures were used to test TmrA production in a variety of conditions; a schematic of the final protocol is shown in Figure 1. To establish necessary conditions for successful expression, TmrA was initially tested with and without *pBAD42-BtuCEDFB* for *btu* operon co-expression (44), and in the presence and absence of oxygen during induction. While TmrA expression was evident in all conditions (Figure S1/SI Text 3), its solubility in each case was difficult to distinguish as very little enzyme was visible and overlapped with the host’s proteins. For this reason, an activity assay, specifically the level of chloroform reduction observed from soluble lysate fraction, was used to indicate the relative production of active TmrA under each set of conditions. All assays were performed under anaerobic conditions using titanium(III) citrate as the reductant, and methyl viologen as the electron donor. The assay results were measured by end-point detection after 18-24 hr using GC-FID to quantify the substrate and product(s). We found that the co-expression of the *btu* operon paired with TmrA induction under anaerobic conditions was essential to produce active TmrA in *E. coli* (Figure 2A). Under these two conditions 76 ± 4 nmol of dichloromethane (DCM) was produced (significantly higher than the limit of detection of 5 nmol for DCM on this instrument), whereas with no Btu co-expression or in the presence of oxygen no DCM was detected.

**Figure 2.**
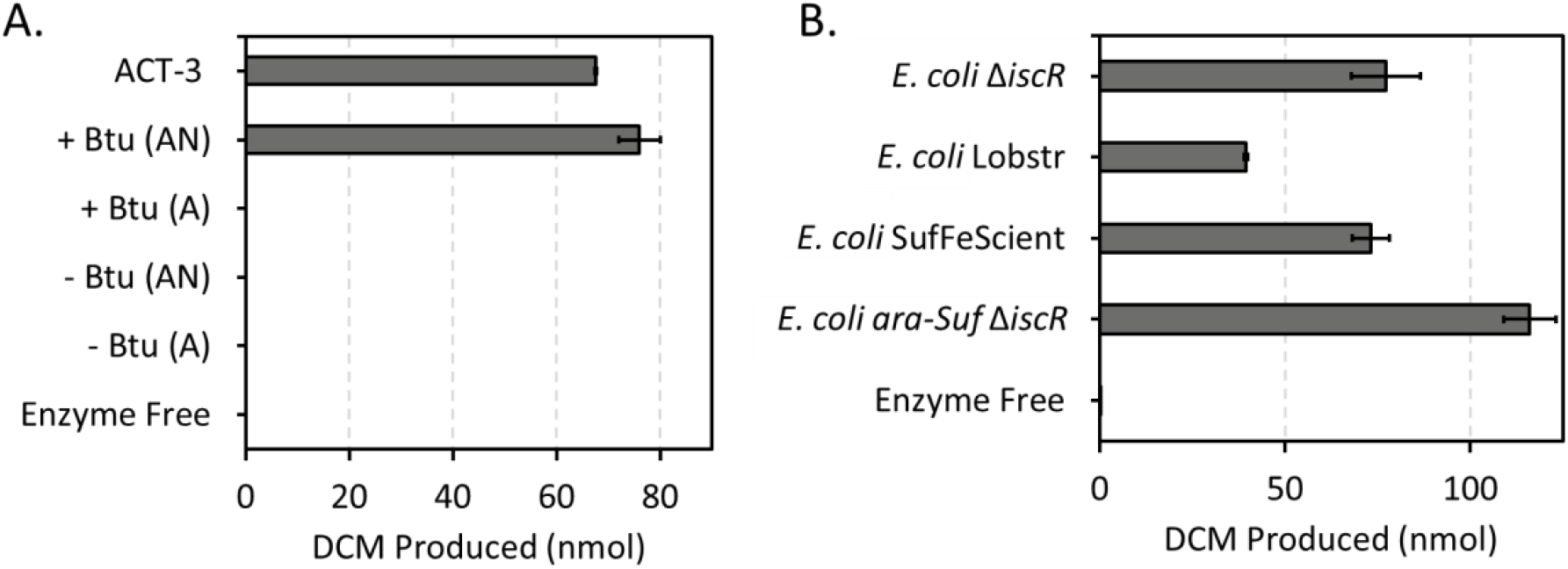
Reduction of chloroform to dichloromethane (DCM) by *E. coli* cell-free extracts expressing TmrA over a period of 24 hr. (A) Comparison of TmrA expressed in *E. coli* Δ*iscR* with (+ Btu) and without (− Btu) *pBAD42-BtuCEDFB* plasmid expression, and with induction under anaerobic (AN) or aerobic (A) conditions. (B) Comparison of TmrA expressed in different *E. coli* strains with *pBAD42-BtuCEDFB* co-expression and under anaerobic conditions. ACT-3 mixed culture cell-free extract was used as a positive control and an enzyme free negative control was used. Error bars are standard deviation between replicates (n = 6, except for ACT-3, enzyme free, and aerobic samples where n = 3).

A variety of additional conditions were tested to see if the observed activity could be enhanced, these included: varying lysis buffer detergent, inclusion of the TAT signal peptide, and testing different *E. coli* strains. The detergents tested were 0.1% Triton X-100, 1% CHAPS, and 1% digitonin; there was only a minor difference in DCM production between these detergents (82 ± 1, Triton X-100; 76 ± 2, digitonin; 67.9 ± 0.3, CHAPS), so all subsequent experiments were continued with Triton X-100. However, this flexibility may not be true for all RDases as some *Dehalococcoides* RDases were shown to be inhibited by certain detergents (59). The inclusion of the TAT sequence when expressing TmrA also did not have a notable effect on the enzyme activity; although, it could interfere with downstream applications, so we chose to use the truncated TmrA. Finally, we tested three different strains of *E. coli* that have enhanced [4Fe-4S] production: *E. coli* Δ*iscR* (49, 50), *E. coli* SufFeScient (51), and *E. coli ara-Suf* Δ*iscR*, as well as one typical expression strain, *E. coli* BL21(DE3) Lobstr. Lysates from *E. coli* Δ*iscR* and SufFeScient had similar levels of chloroform reduction, whereas there was a 30% increase in activity observed from TmrA expressed in *E. coli ara-Suf* Δ*iscR* (Figure 2B). All the [4Fe-4S] strains displayed at least double the activity as the typical strain *E. coli* Lobstr. *E. coli ara-Suf* Δ*iscR* was subsequently used as the primary expression strain, unless the RDase of interested displayed poor expression in this strain, in which case *E. coli* Δ*iscR* was used.

The implementation of the *pBAD42-BtuCEDFB* plasmid to increase the uptake of cobalamin was key and essential to successful expression. It is now well known that a cobamide co-factor is critical to protein folding and activity in RDases. The requirement of a cobamide during protein expression, in previous studies and this work, and its buried position within the known crystal structures suggests that the cobamide is needed for the enzyme to fold into its catalytically active form. Induction under anaerobic conditions was also necessary, most likely to prevent oxidative damage to the [4Fe-4S] clusters, but perhaps the addition of a strong reductant into the medium could allow for aerobic induction. The use of *E. coli* strains with increased [4Fe-4S] cluster production or stability also improved the level of activity seen in the lysates compared to conventional strains (Figure 2B). These clusters are essential for electron transport within the enzyme. The specific strain of *E. coli* used for [4Fe-4S] production may depend on the desired RDase as the expression levels of some enzymes varied depending on the strain. Overall, this expression system successfully reached the goal of producing active respiratory RDases in *E. coli*. Furthermore, this expression system was successful on an RDase from the obligate OHRB *Dehalobacter* and may be particularly useful for studying the organohalide respiration process.

The only other account of *E. coli* being used for the production of a functional RDase, without cofactor reconstitution, was the expression of NpRdhA (48). Similarly, the authors improved cobalamin transport to support expression and cofactor incorporation (48). However, NpRdhA, being a non-respiratory metabolic (i.e., not membrane associated) RDase which acts on relatively soluble halogenated phenols and hydroxybenzoic acids, is itself more soluble and more oxygen tolerant than respiratory RDases. NpRdhA was expressed in a conventional *E. coli* strain both with and without the co-expression of BtuB from the *btu* operon (48). The addition of the BtuB co-expression increased the cobalamin occupancy by 9% and doubled the enzyme yield (48), but was not a determining factor in the production of active protein, contrary to what we observe from the respiratory RDases. NpRdhA is not a suitable comparator for the systems presented here.

### Comparing heterologous RDase substrate specificity

We applied the optimized expression system to a series of other enzymes to validate its reproducibility. CfrA and DcrA, chloroethane reductases from *Dehalobacter* sp. CF and DCA, respectively, in the ACT-3 mixed culture, were chosen to be tested as they have >95% amino acid sequence identity to TmrA but have displayed differing substrate preferences when isolated and tested from their native producers (14, 27). CfrA was experimentally shown to reduce chloroform and 1,1,1-trichloroethane (1,1,1-TCA) but not 1,1-DCA; DcrA accepts 1,1-dichloroethane (1,1-DCA) as a substrate, but not 1,1,1-TCA or CF; and TmrA accepts all three substrates (14, 27). As well, 1,1,2-trichloroethane (1,1,2-TCA) is a substrate for all three enzymes, although the products of reaction vary as 1,1,2-TCA can undergo both dihaloelimination or hydrogenolysis (14, 60). DcrA has not been explicitly assayed against 1,1,2-TCA, but the organism that it comes from can use 1,1,2-TCA as an electron acceptor so it has assumed activity on this substrate (60). The known distinction between these enzymes’ substrate ranges made them good subjects to confirm that the expressed enzymes exhibit the same specificity. Both CfrA and DcrA were successfully expressed in small-scale cultures and the resulting cell lysates displayed activity on their expected substrates.

TmrA, CfrA, and DcrA were all assayed against several chloroalkanes: chloroform, 1,1,1-TCA, 1,1,2-TCA, and 1,1-DCA. The amount (nmol) of each dechlorinated product produced in the assays are given in Table 1, some substrates yielded multiple products depending on the reaction pathway. Heat deactivated controls were used for each substrate, none of which showed notable reduction of the substrate except 1,1,1-TCA to 1,1-DCE though this transformation was also seen in the enzyme free control (Table S8).While the specific activity of each enzyme cannot be compared from the lysate assays as the amount of RDase is unknown, their substrate preferences and products can be compared. TmrA and CfrA both reduced chloroform, 1,1,1-TCA, and 1,1,2-TCA via hydrogenolysis; although, TmrA transformed a considerable amount of 1,1,2-TCA into the β-elimination product, vinyl chloride (VC), while CfrA only produced a minute amount of VC. TmrA was also able to reduce 1,1-DCA to chloroethane (CA) while CfrA had negligible activity on this substrate consistent with previous studies (27). In contrast, DcrA reduced 1,1-DCA, but was not active on CF and barely on 1,1,1-TCA (Table 1), also consistent with previous studies (27). The one surprising result was the high production of 1,1-DCE from 1,1,1-TCA as this is only a minor by-product in the cultures and is mainly thought of as an abiotic reaction (61). Since 1,1-DCE is also seen in the negative controls we anticipate it to be an abiotic reaction accelerated by a component of the buffer. The observed substrate selectivity, particularly those of CfrA and DcrA, act as validation that the heterologously expressed enzymes behave similarly to those partially purified and characterized from the native organisms.

**Table 1.** Reduction of chlorinate substrates over 24 hr by the cell extracts of *E. coli* expressing TmrA, CfrA, and DcrA. All assays were done with approximately 4-10 µg of total protein. Error is the standard deviation between replicates, number of replicates are indicated for each sample.

|  |  | Enzyme-Free | TmrA | CfrA | DcrA |
| --- | --- | --- | --- | --- | --- |
| Known Substrates |  |  | CF; 1,1,1-TCA; 1,1,2-TCA; 1,1-DCA | CF; 1,1,1-TCA; 1,1,2-TCA | 1,1-DCA; 1,1,2-TCA |
| Substrate | Product(s) | Amount Produced (nmol) |  |  |  |
| CF | DCM | n.d. <sup>a</sup> | 101 ± 7 <sup>b</sup> | 90 ± 10 <sup>b</sup> | n.d. <sup>a</sup> |
| 1,1,1-TCA | 1,1-DCA | 0.09 ± 0.06 <sup>a</sup> | 7 ± 3 <sup>b</sup> | 13 ± 1 <sup>b</sup> | 0.20 ± 0.02 <sup>a</sup> |
|  | 1,1-DCE* | 1.3 ± 0.3 <sup>a</sup> | 12.4 ± 0.8 <sup>b</sup> | 11.7 ± 0.2 <sup>b</sup> | 12.1 ± 0.1 <sup>a</sup> |
| 1,1,2-TCA | 1,2-DCA | n.d. <sup>a</sup> | 38 ± 6 <sup>c</sup> | 35 ± 5 <sup>c</sup> | n.d. <sup>a</sup> |
|  | VC* | n.d. <sup>a</sup> | 11 ± 2 <sup>c</sup> | 0.4 ± 0.2 <sup>c</sup> | 1.0 ± 0.2 <sup>a</sup> |
| 1,1-DCA | CA | n.d. <sup>a</sup> | 9 ± 5 <sup>c</sup> | 0.07 ± 0.05 <sup>c</sup> | 13 ± 4 <sup>b</sup> |
|  | VC* | n.d. <sup>a</sup> | n.d. <sup>c</sup> | n.d. <sup>c</sup> | 0.04 ± 0.03 <sup>b</sup> |

CF = chloroform, DCM = dichloromethane, TCA = trichloroethane, DCA = dichloroethane, VC = vinyl chloride, CA = chloroethane, n.d. = not detected
\* β-elimination pathway, <sup>a</sup> n = 2, <sup>b</sup> n = 6, <sup>c</sup> n = 4

The successful expression of CfrA and DcrA was expected after optimizing the system on TmrA since the enzyme sequences are almost identical to each other and are thus likely to behave similarly. While these three enzymes have been previously studied, it was still intriguing to remove any other biological factors from the native organisms and confirm their unique substrate preferences in a heterologous system. In agreement with the literature, CfrA was much more efficient at reducing trichloroalkanes, whereas DcrA can almost exclusively reduce 1,1-DCA (27). TmrA displays activity against all the chloroalkanes assayed, and reduced 1,1,2-TCA to both 1,2-DCA and VC. The observed activity differences by each of these enzymes confirms that it is their amino acid sequence that affects their activity and not external factors such as the type of cobamide, protein-protein interactions, or other cellular components.

### TmrA purification and specific activity

TmrA was elected for large scale purification trials because TmrA has published activity and expression data to compare against. TmrA was semi-purified from a 1 L expression culture using nickel affinity chromatography. An SDS-PAGE of the TmrA purification is shown in Figure S2A/SI Text 3. The concentrated protein sample was a yellow-brown colour indicating the presence of [4Fe-4S] clusters. TmrA was purified to an approximate purity of 50%, 2.2 ± 0.2 mg of protein was obtained from about 7.5 g wet cell mass. For details about the purification and purity estimation see Figure S2B and Table S6 of SI Text 3. To further assess the quality of the purification, the cofactors were extracted and quantified to measure their occupancy ratio. RDases are expected to have eight iron atoms from two [4Fe-4S] clusters. After adjusting for purity (i.e., assume the cofactors were only coming from TmrA), TmrA was found to have 6.2 ± 0.6 mol iron per mol enzyme. The cobalamin occupancy was measured by two methods; the first was extracted into water and measured by absorbance at 367 nm, characteristic of dicyanocobalamin. Using this method and correcting for purity, TmrA was estimated to have cobalamin occupancies of 62 ± 13% for TmrA. However, using the second method which involved extraction into acidified methanol and quantification by UHPLC-MS and correcting for purity, the enzymes occupancies were measured to be 24 ± 2%. Finally, we found that TmrA was able to withstand freeze-thaw cycles without loss of activity and was able to retain all activity after being exposed to O_2_ for 3 hr with 1 mM TCEP in the buffer (longer periods have not been tested; data not shown).

The specific activity of TmrA was determined for several substrates; specific activity was reported in nkat (nmol product/s) per mg protein. TmrA activity was measured for all of its known substrates that showed activity in the lysate assays. Assays with the purified enzyme also identified other transformations that were not observed in lysate, such as 1,2-DCA to CA and VC, as well as 1,1-DCA to VC. TmrA had the highest specific activity towards chloroform and 1,1,2-TCA into both 1,2-DCA and VC, whereas it had low specific activity towards 1,1,1-TCA and the less chlorinated substrates 1,1-DCA and 1,2-DCA (Figure 3). When concentrated, TmrA was observed to reduce the dichloroethanes and perform both hydrogenolysis and β-elimination reactions on all chloroethane substrates. The reduction of 1,1,1-TCA to 1,1-DCE is not shown as there were similar concentrations in the enzyme-free negative controls so the biotic activity could not be distinguished (Table S11). There was also production of 1,1-DCE in the lysate samples, though the production from the enzymes was 10-fold higher than the enzyme-free control. In the purified protein case, the production levels were equivalent to the negative control which highly suggests it is not a biotic reaction.

**Figure 3.**
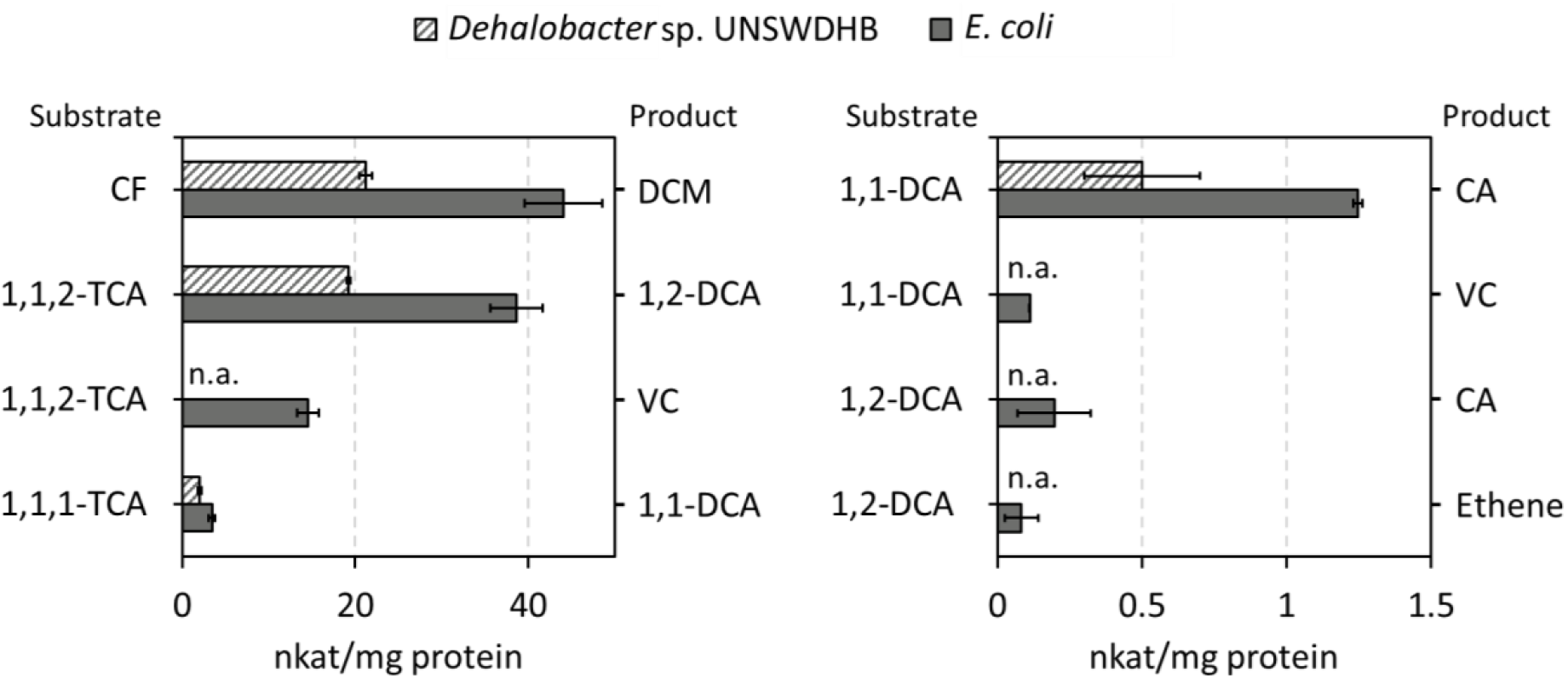
Specific activity of TmrA for substrate-product transformations, where nkat is defined as the rate of product formation in nmol/s. Data for TmrA purified from the endogenous host *Dehalobacter* sp. UNSWDHB (striped bars) (14), and TmrA heterologously expressed and purified from *E. coli* (solid bars) are shown. Substrates are indicated on the left axis label and product is indicated on the right axis label. Error bars are from reported standard deviation for the literature values, and the standard deviation of purified enzyme assays (n = 3). n.a. = not available.

Not only did the substrate profiles match the literature, the purified TmrA had a specific activity that corresponded well to the native enzyme purified from *Dehalobacter* sp. UNSWDHB. The native enzyme was found to be most active on CF and 1,1,2-TCA, and its specific activity towards its substrates was within the same magnitude of the heterologously expressed TmrA (Figure 3). This consistency between the enzymes confirms that the heterologous system is representative of the native enzymes.

The use of *E. coli* as the heterologous host has several advantages over the hosts previously used for respiratory RDases, namely *S. blattae* and *B. megaterium*. One advantage is higher protein yields in *E. coli*. From 1 L of culture, we were able to obtain approximately 2 mg of TmrA whereas to get 0.9 mg of TmrA at a similar purity, 15 L of *B. megaterium* was required (35). Furthermore, the RDase expression vectors can be commercially produced in typical *pET* vectors while those for *B. megaterium* and *S. blattae* generally have to be manually cloned (33, 35, 36). The cofactor occupancy of the RDases from this *E. coli* expression system are also comparable to other heterologous hosts. The molar ratio of iron to enzyme was reported to be 7.26 ± 0.48 mol iron/mol TmrA produced in *B. megaterium* and we measured 6.2 ± 0.6 mol iron/mol TmrA from our system (35). TmrA from *B. megaterium* was calculated to have 52 ± 3% cobalamin occupancy (35), using a similar absorbance based method we measured an occupancy of 62 ± 13%. Although, the variation in cobalamin occupancy measured between the two methods used in this study suggests that the true occupancy is within 20-60%. Our system seems to perform similarly to those using alternative heterologous hosts, is reproducible, and facilitates high-throughput protein production.

### Expression and purification of a lindane reductase

HchA, identified from *Dehalobacter* sp. HCH1 as a lindane or γ-hexachlorocyclohexane (γ-HCH) reductase (62), was chosen to test the generalizability of the expression system because it has low amino acid sequence identity (32.8%) to TmrA. This enzyme was recently identified using BN-PAGE in enrichment cultures that dechlorinate HCH via three sequential dechlorination steps to form monochlorobenzene (MCB) and benzene (62, 63). HchA was successfully expressed and was assayed on several of the chlorinated solvents as well as on two isomers of HCH (γ-HCH and α-HCH).

HchA produced primarily monochlorobenzene (MCB) from its known substrate γ-HCH (Table 2). Interestingly, it transformed more of the α-HCH isomer and to a higher proportion of benzene. However, there was significant reduction of both isomers by the HchA heat-killed controls and by free cobalamin (Table 2), so the biotic activity needed to be further confirmed (see below). HchA was also able to take 1,1,2-TCA as a substrate and transform it to VC to a greater extent than the cobalamin negative control. Some highly chlorinated substrates, such as γ-HCH and 1,1,2-TCA, are more susceptible to abiotic reduction, whereas others including chloroform are much more resistant to reduction. HchA readily dechlorinated these permissive substrates and primarily seems to catalyze dihaloelimination reactions.

**Table 2.** Reduction of chlorinate substrates over 24 hr by the cell extracts of *E. coli* expressing HchA and the relevant negative controls. All assays were done with approximately 4-10 µg of total protein, or 100 nmol of hydroxycobalamin. Error is standard deviation of replicates; number of replicates is indicated for each sample.

| Substrate | Product(s) | HchA | Heat Killed HchA | Cobalamin | Catalyst Free |
| --- | --- | --- | --- | --- | --- |
|  |  | Amount Produced (nmol) |  |  |  |
| CF | DCM | n.d. <sup>a</sup> | n.t. | n.t. | n.d. <sup>a</sup> |
| 1,1,2-TCA | 1,2-DCA | n.d. <sup>a</sup> | n.t. | n.d. | n.d. <sup>a</sup> |
| | VC* | 18 ± 6 <sup>a</sup> | n.t. | 9.5 | n.d. <sup>a</sup> |
| 1,1-DCA | CA | n.d. <sup>a</sup> | n.t. | n.t. | n.d. <sup>a</sup> |
| γ-HCH | MCB | 7 ± 1 <sup>b</sup> | 7.3 ± 0.7 <sup>c</sup> | 9 ± 2 <sup>a</sup> | 0.4 ± 0.1 <sup>c</sup> |
| | Benzene | 0.07 ± 0.07 <sup>b</sup> | 0.12 ± 0.07 <sup>c</sup> | 0.21 ± 0.04 <sup>a</sup> | 0.07 ± 0.05 <sup>c</sup> |
| α-HCH | MCB | 18 ± 1 <sup>a</sup> | n.t. | 5.1 | 0.58 |
| | Benzene | 0.6 ± 0.1 <sup>a</sup> | n.t. | 0.15 | n.d. |

CF = chloroform, DCM = dichloromethane, TCA = trichloroethane, DCA = dichloroethane, VC = vinyl chloride, CA = chloroethane, MCB = monochlorobenzene, n.d. = not detected, n.t. = not tested
\* Dihaloeelimination pathway, <sup>a</sup> n = 2, <sup>b</sup> n = 6, <sup>c</sup> n = 3.

To probe relative importance of biotic and abiotic reduction of γ-HCH, HchA was scaled up and purified using the same methods as TmrA. HchA was enriched to 39% purity and 2.7 ± 0.4 mg of protein was obtained from about 7 g wet cell mass. For details about the purification see Table S6/SI Text 3. The cofactors were also quantified to estimate their occupancy. HchA was found to have 6.4 ± 0.8 mol iron per mol enzyme, a cobalamin occupancy of 76 ± 14% using the UV-Vis method, and a cobalamin occupancy of 22.1 ± 0.7% using the LC-MS method. These numbers are consistent with what was observed for TmrA. HchA was also found to be tolerant of freeze-thaw cycles and retained all activity after 30 min of O_2_ exposure (with 1 mM TCEP), but only held 60 ± 35% of its activity after 1 hr of exposure (data not shown).

HchA specific activity was measured on γ-HCH, 1,1,2-TCA, and 1,1,1-TCA. HchA had the highest level of activity transforming 1,1,2-TCA to VC, whereas it showed minor transformation of 1,1,2-TCA to 1,2-DCA (0.02 ± 0.01 nkat/mg) and 1,1,1-TCA to 1,1-DCA (0.11 ± 0.01 nkat/mg; Figure 4A). The activity on its primary substrate γ-HCH to MCB was 1.7 ± 0.4 nkat/mg, the transformation to benzene was not shown as it was also produced in the negative controls. Since the reduction of γ-HCH can also occur abiotically by free cobalamin, the rate of reduction was compared when normalized by mol of catalyst (i.e., HchA and cobalamin). HchA was approximately fourteen times faster than the abiotic reaction (Figure 4B). Further, the killed control of HchA no longer had significant activity indicating that the activity seen in the lysates was most likely due to free cobalamin and longer incubation periods. These results suggests that though γ-HCH is reduced by free cobalamin, the enzyme is clearly more effective.

**Figure 4.**
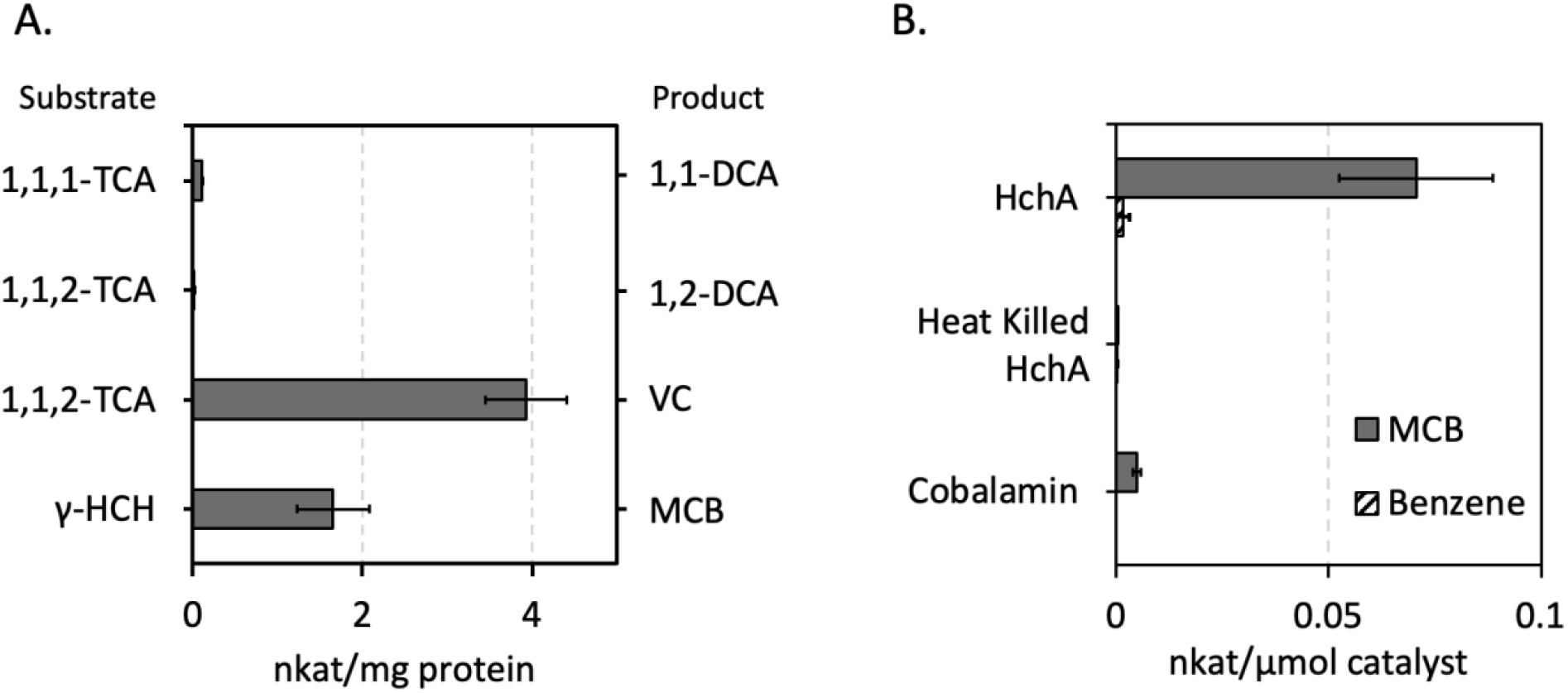
(A) Specific activity of HchA for substrate-product transformations. (B) The rate of monochlorobenzene (MCB) and benzene production from purified HchA, heat killed HchA, and free cobalamin (same molarity of catalysts were used). The nkat is defined as the rate of product formation in nmol/s, this rate is normalized to mols of catalyst. Error bars are the standard deviation between assays (n = 3).

Further, HchA displayed activity that was not tested previously, most likely due to the limited enzyme available from BN-PAGE separation. HchA was only ever documented to reduce γ-HCH though it seems to be more effective at transforming the alpha isomer and the permissive substrate, 1,1,2-TCA. These reactions seen by HchA in the lysates are all expected to be dihaloelimination reactions, but when HchA was concentrated after enrichment it was also observed to have slight hydrogenolysis activity against the 1,1,1-TCA. TmrA was tested against γ-HCH as well but did not catalyze any meaningful transformations (data not shown), suggesting that HchA has some specialization for HCH. HchA poses an interesting comparison to other RDases that predominantly undergo hydrogenolysis reactions to understand how the enzymes distinguish between reaction pathways.

### Expression of putative *rdhA* genes

To further test the application of the expression system, we expressed two putative RdhAs, DHB14 and DHB15, that had been previously cloned from the genomic DNA of *Dehalobacter* sp. strains CF and DCA, respectively. These enzymes cluster to a group of *Dehalobacter* and *Desulfitobacterium* RDases including several PceA and TcbA enzymes (Figure 5A), which accept unsaturated and aromatic substrates (29, 31, 64). Based on the similarities to known chloroethene reductases, DHB14 and DHB15 were expressed, the cell lysates were assayed against perchloroethene (PCE) and trichloroethene (TCE), and the presence of dechlorinated products was analyzed (Figure 5B). DHB15 reduced PCE to TCE and a small amount of *cis*-dichloroethene (cDCE), DHB14 only reduced a minor amount of PCE to TCE. Both enzymes reduced a modest amount of TCE to cDCE. The enzymes were also tested for activity on 1,2-dichloroethane (1,2-DCA); however, no product was detected after 24 hr of incubation. The activity of these two enzymes demonstrate that this expression system has potential to functionally characterize the numerous *rdhA* genes encoded in *Dehalobacter* spp. genomes.

**Figure 5.**
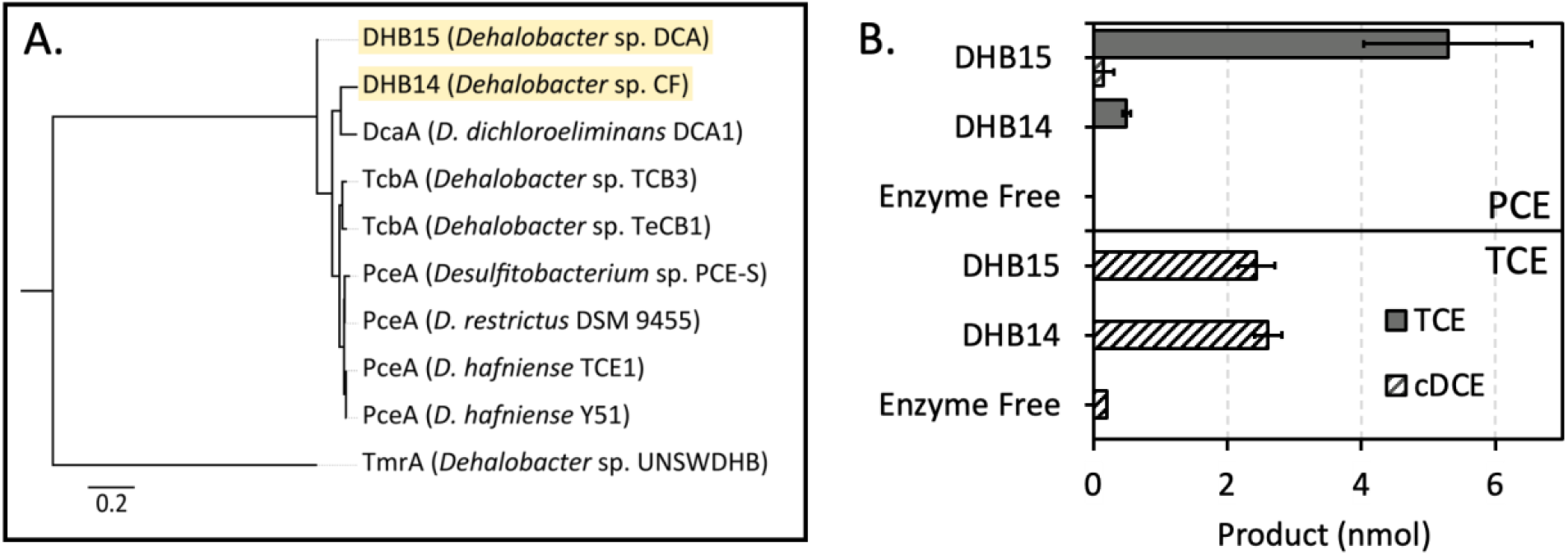
(A) Maximum-likelihood (ML) tree of DHB14 and DHB15 RdhA amino acid sequences to show relation to the closest characterized RDases in comparison to TmrA. (B) Reduction of either PCE (top) or TCE (bottom) by DHB14 and DHB15 in *E. coli* cell-free extract, assay was done over 24 hr. Error bars are the standard deviation between replicates (n = 4), except negative controls (n = 2). An amino acid alignment was built using the Geneious v8.1.9 MUSCLE plugin, and the ML tree was constructed using RAxML v8.2.12 Gamma GTR protein method with 100 bootstraps, the highest scoring tree was selected. Scale indicates average number of substitutions per amino acid site.

### *Dehalococcoides* RDase expression attempts

The heterologous expression of several RDases from *Dehalococcoides* spp. was attempted using this expression system. Three characterized chloroethene reductases were cloned and expressed: VcrA from *D. mccartyi* VC, BvcA from *D. mccartyi* BAV1, and TceA from *D. mccartyi* 195. Expression of these enzymes was apparent by SDS-PAGE; however, no activity was ever detected from the expression culture lysates. The details of all conditions tested for the expression of these RDases can be found in the Supplemental Information Text 4.

The inability to achieve activity from the *Dehalococcoides* spp. enzymes could be due to the absence of an unknown component required for folding such as an organism-specific chaperone protein. Further work is needed to test more expression conditions for RDases from the Chloroflexi *Dehalococcoides* and *Dehalogenimonas* to identify the essential factors in their production. Nevertheless, the system we have presented provides an excellent starting point for future efforts.

### Significance and future work

Developing a heterologous system for RDase expression in *E. coli* has been a goal of the dehalogenation community for the past several decades. Here we have presented a solution for the expression of *Dehalobacter* derived RDases that will allow high yield expression and purification, enabling future structural and mechanistic studies. We demonstrated that this system was able to be applied to various RDases from *Dehalobacter* having as little as 28% amino acid sequence identity to each other. Given the high similarity between *Dehalobacter* and *Desulfitobacterium* RDases, we predict that this system would perform similarly for RDases from other Firmicutes.

We anticipate that this expression system will expedite the characterization of the RDase enzyme family and advance the future of enzyme-driven bioremediation. Mutational studies can now be carried out to identify key amino acids required for activity and specificity. The impact of the type of corrinoid incorporated into the RDase on activity and substrate range can also be more definitively investigated. However, the current standard assays used in this work require development of analytical methods and individual standards for each substrate and product to detect activity; a general assay to measure non-specific activity would allow data to be obtained more quickly and would facilitate high-throughput analysis.

This work also has many implications for bioremediation. Individual RDases can be screened on wider substrates panels to better delineate substrate ranges and to quantify effects of putative inhibitors. This knowledge would better inform the choice of bioaugmentation culture(s) when targeting complex contamination sites. Furthermore, this expression system will also allow the functional determination for the huge number of undescribed *rdhA* genes present in metagenomes, which in turn could identify novel candidate organisms for bioremediation. Finally, having an established expression method in *E. coli* gives the opportunity to rationally engineer and evolve RDases to dehalogenate emerging and highly halogenated compounds. However, two bottlenecks that still must be addressed to truly jumpstart the future studies of RDases are 1) generalizing or extending the assay to Chloroflexi, and 2) developing a high-throughput and generalizable assay to make data production quicker and allow for the testing of a wide array of substrates and enzymes.

In conclusion, we demonstrate the necessity of cobalamin and anaerobic conditions for the heterologous expression of respiratory RDases from obligate OHRB. The implementation of the *pBAD42-BtuCEDFB* expression vector allowed for the active expression of RDases in *E. coli*, making this the first report of respiratory RDases actively expressed from *E. coli*. We demonstrated that this system is easy to modify and was widely applicable within the *Dehalobacter* genus by expressing and obtaining activity from six unique RDases. The expressed RDases were able to be purified at high yields making this system useful for future analyses that require a lot of material. We expect that this expression system will allow for the study and characterization of many RDases that have so far remained elusive.

## ACKNOWLEDGEMENTS

This study was supported by the National Sciences and Engineering Research Council of Canada (NSERC PGS-D to KJP) and a Canadian Research Chair to EAE.

EAE and KJP conceived of the experiments. RF performed the LC-MS separation and analysis of cobalamin, and KJP conducted all other experiments. KJP and EAE drafted the manuscript.

We would like to thank the Booker Lab from Pennsylvania State University for gifting the *pBAD42-BtuCEDFB* plasmid, and the Antony Lab from the St. Louis University School of Medicine and Kiley lab from the University of Wisconsin-Madison for gifting us the *E. coli* SufFeScient and *ara-Suf* Δ*iscR* strains. We would like to thank Connor Bowers for cloning and testing the activity of the SUMO fusion constructs. We would like to thank Anna Khusnutdinova for helpful advice in working with anaerobic enzymes and more generally the Yakunin and Savchenko labs at the University of Toronto for sharing expression vectors and several RDase constructs, and for their support during this study. Finally, we would like to thank Prof. Patrick Hellenbeck for providing the *E. coli* Δ*iscR* strain.

## Notes

### Competing Interest Statement

The authors have declared no competing interest.

