## Supplemental Information for "Heterologous expression of *Dehalobacter* spp. respiratory reductive dehalogenases in *Escherichia coli*"

#### SI Text 1 – Partially Characterized RDases

This SI section include two tables, one describing all partially characterized RDases we could find and their characterization methods, and another describing published attempts at heterologous expression. Most of the described enzymes are described in the Reductive Dehalogenase Database (https://rdasedb.biozone.utoronto.ca/) (1).

**Table S1. Characterized reductive dehalogenases.** Bolded organisms are obligate organohalide respiring bacteria.

| Enzyme | Organism*/Culture | Substrate class | Method of Functional Determination | Reference |
|---|---|---|---|---|
| 3,5-DCPh-CprA5 | *Desulfitobacterium* | Chlorophenols | Purification from native producer | (2) |
| 3-CBA RDase | *Desulfomonile tiedje* | Chlorobenzoates | Purification from native producer | (3) |
| BhbA | *Comamonas* sp. 7D-2 | Bromobenzoate | Purification from native producer | (4) |
| BvcA | *Dehalococcoides*/ KB-1 consortium | Chloroethenes | BN-PAGE | (5) |
| BvcA | ***Dehalococcoides mccartyi* BAV1** | Chloroethenes | Transcriptomics, BN-PAGE | (6, 7) |
| CbrA | ***Dehalococcoides mccartyi* CBDB1** | Chlorobenzenes | BN-PAGE | (8) |
| CerA | ***Dehalogenimonas* sp. GP** | Chloroethenes | Proteomics | (9) |
| CfrA | ***Dehalobacter* sp. CF** | Chloroethanes | BN-PAGE | (10) |
| Cl-OHPA CprA | *Desulfitobacterium hafniense* DCB-2 | Chlorohydroxy-phenylacetate | Purification from native producer | (11) |
| CprA | *Desulfitobacterium* sp. Viet-1 | Chlorohydroxy-phenylacetate | Metagenomics | (12) |
| CprA | *Desulfitobacterium* sp. KBC1 | Chlorophenols | Purification from native producer | (13) |
| CprA | *Desulfitobacterium* sp. PCE-1 | Chlorophenols | Purification from native producer | (14) |
| CprA | *Desulfitobacterium chlororespirans* | Chlorophenols | Purification from native producer | (15) |

| | | | | |
|---|---|---|---|---|
| CrdA | *Desulfitobacterium hafniense* PCP-1 | Chlorophenols | Purification from native producer | (16) |
| CtrA | *Desulfitobacterium* sp. PR | Chloroethanes | Proteomics | (17) |
| DcaA | *Desulfitobacterium dichloroeliminans* DCA1 | Chloroethanes | Heterologous expression | (18) |
| DcpA | ***Dehalogenimonas lykanthroporepellans*** | Chloropropanes | BN-PAGE | (19) |
| DcrA | ***Dehalobacter* sp. DCA** | Chloroethanes | BN-PAGE | (10) |
| DebcprA | ***Dehalobacter* sp. TCP1** | Chlorophenols | Transcriptomics | (20) |
| GPceA | *Geobacter* sp. | Chloroethenes | Heterologous expression | (21) |
| GeobRD | *Geobacter* sp./ KB-1 consortium | Chloroethenes | BN-PAGE | (5, 6) |
| HchA | ***Dehalobacter* sp. HCH1** | Hexachloro-cyclohexane | BN-PAGE | (22) |
| MbrA | ***Dehalococcoides* sp. MB** | Chloroethenes | BN-PAGE | (23) |
| NpRdhA ‡ | *Nitratireductor pacificus* pht-3B | Halophenols, Halohydroxy-benzoates | Heterologous expression; 3D structure obtained | (24, 25) |
| PcbA1 | ***Dehalococcoides* sp. PCB1** | Polychlorinated biphenyls | Metagenomics, Cell extract assay | (26) |
| PcbA4 | ***Dehalococcoides* sp. PCB4** | Polychlorinated biphenyls | Metagenomics, Cell extract assay | (26) |
| PcbA5 | ***Dehalococcoides* sp. PCB5** | Polychlorinated biphenyls | Metagenomics, Cell extract assay | (26) |
| PceA | ***Dehalococcoides mccartyi* 195** | Chloroethenes | Purification from the native producer | (27) |
| PceA | *Desulfitobacterium* sp. PCE-1 | Chloroethenes | Purification from the native producer | (14) |
| PceA | *Shewanella sediminis* | Chloroethenes | Transcriptomics, Cell extract assay | (28) |
| PceA | *Desulfitobacterium* sp. PCE-S | Chloroethenes | Purification from the native producer | (29) |
| PceA | ***Dehalobacter restrictus* DSM 9455** | Chloroethenes | Purification from the native producer | (30) |
| PceA | *Desulfitobacterium hafniense* TCE1 | Chloroethenes | Purification from the native producer | (14) |

| | | | | |
|---|---|---|---|---|
| PceA | *Desulfitobacterium hafniense* Y51 | Chloroethenes | Heterologous expression | (31) |
| PceA | *Sulfurospirillum multivorans* N | Chloroethenes | Transcriptomics, Cell extract assay | (32) |
| PceA ‡ | *Sulfurospirillum multivorans* 12446 | Chloroethenes | Purification from the native producer with genetic modification; 3D structure obtained | (33) |
| PceA-DCE | *Sulfurospirillum* mixed culture SL2 | Chloroethenes | Transcriptomics, Cell extract assay | (32) |
| PceA-TCE | *Sulfurospirillum* mixed culture SL2 | Chloroethenes | Transcriptomics, Cell extract assay | (32) |
| PentaCPh-CprA3 | *Desulfitobacterium hafniense* PCP-1 | Chlorophenols | Purification from the native producer | (34) |
| PrdA | *Desulfitobacterium* sp. KBC1 | Chloroethenes | Transcriptomics, Cell extract assay | (35) |
| RdhA3 | *Desulfitobacterium hafniense* DCB-2 | Chloroethenes | Heterologous expression | (31) |
| TcbA | ***Dehalobacter* sp. TCB3** | Chlorobenzenes | BN-PAGE | (22) |
| TcbA | ***Dehalobacter* sp. TeCB1** | Chlorobenzenes | BN-PAGE | (36) |
| TceA | ***Dehalococcoides*/ KB-1 consortium** | Chloroethenes | BN-PAGE | (5) |
| TceA | ***Dehalococcoides mccartyi* 195** | Chloroethenes | Purification from the native producer | (27) |
| TdrA | ***Dehalogenimonas* sp. WBC-2** | Chloroethenes | BN-PAGE | (37) |
| ThmA | ***Dehalobacter* sp. PR** | Chloroethanes | BN-PAGE | (38) |
| TmrA | ***Dehalobacter* sp. UNSWDHB** | Chloroethanes | Purification from the native producer, Heterologous expression | (39, 40) |
| VcrA | ***Dehalococcoides*/ KB-1 consortium** | Chloroethenes | BN-PAGE | (5) |
| VcrA | ***Dehalococcoides* sp. VS** | Chloroethenes | Purification from the native producer, Heterologous expression | (41, 42) |
| VcrA | ***Dehalococcoides* sp. WBC-2** | Chloroethenes | BN-PAGE | (37) |

* Organism may be isolate or identified in mixed culture by metagenomics
‡ Enzyme has had crystal structure solved
BN-PAGE = Blue native - PAGE

**Table S2. Heterologously expressed reductive dehalogenases.** Bolded organisms are obligate organohalide respiring bacteria, enzymes in red had no activity detected on the substrates tested.

| Enzyme | Native Organism | Heterologous Host | Substrates Tested | Additional Details | Reference |
|---|---|---|---|---|---|
| PceA | *Sulfurospirillum multivorans* | *E. coli* | PCE | Expression was seen but no activity detected. | (43) |
| PceA | *Desulfitobacterium hafniense* Y51 | *E. coli* | PCE, | Expression was seen but no activity detected. | (44) |
| PceA | ***Dehalobacter restrictus* DSM 9455** | *E. coli* | N/A | Protein was refolded to incorporate cofactors; no activity detected. | (45) |
| RdhA3 | *Desulfitobacterium hafniense* DCB-2 | *Shimwellia blattae* | PCE, DCPs | Unable to purify using Strep tag. | (31) |
| PceA | *Desulfitobacterium hafniense* Y51 | *Shimwellia blattae* | PCE, TCE, DBA, 1,1,2,2-TeCA | Unable to purify using Strep tag. | (18, 31) |
| VcrA | ***Dehalococcoides* sp. VS** | *E. coli* | VC, 1,1-DCE, 1,2-DCA | Protein was refolded to incorporate cofactors; activity detected. | (42) |
| NpRdhA ‡ | *Nitratireductor pacificus* pht-3B | *Bacillus megaterium, E. coli* | DBP, DBHBA | Enzyme has been actively expressed in multiple hosts and is able to be purified using His tag. 3D Structure obtained | (24, 25) |
| DcaA | *Desulfitobacterium dichloroeliminans* DCA1 | *Shimwellia blattae* | 1,2-DCA, 1,2-DBA, 1,1,2-TCA, 1,1,2,2-TeCA | Protein was co-expressed with RdhT chaperone and enriched using Strep tag. | (18) |
| TmrA | ***Dehalobacter* sp. UNSWDHB** | *Bacillus megaterium* | CF | Purified by nickel affinity and anion exchange chromatography. | (40) |
| PceA | *Geobacter* sp. | *E. coli* | PCE, TCE | Protein was refolded to incorporate cofactors; activity detected. | (21) |

‡ Enzyme has had crystal structure solved; PCE = perchloroethene; TCE = trichloroethene; DCE = dichloroethane; VC = vinyl chloride; TeCA = tetrachloroethane; DCA = dichloroethane DCP = dichlorophenol; DBA = dibromoethane; DBP = dibromophenol; DBHBA = dibromohydroxybenzoic acid; CF = chloroform

### SI Text 2 – Cloning and Expression Information

#### RDase Amino Acid and Nucleotide Sequences

Below are the amino acid sequences and nucleotide sequences of all proteins expressed, predicted TAT sequences that were removed are bolded and underlined, predicted TAT sequences that were not removed for expression are only underlined. The nucleotide sequences are shown as ordered and indicated if codon optimized; if the TAT sequence was not included in the synthetic gene it is omitted in the displayed sequence.

> TmrA | *Dehalobacter* sp. UNSWDHB amino acid

**MDKEKSNNDKPATKINRRQFLKFGAGASSGIAIATAATALGGKSL**IDPKQVYAGTV
KELDELPFNIPADYKPFTNQRNIFGQAVLGVPEPLALVERFDEVRWNGWQTDGSPGLTV
LDGAAARASFAVDYYFNGENSACRANKGFFEWHPKVPELNFKWGDPERNIHSPGVKSA
EEGTMAVKRMARFFGAAKAGIAPFDKRWVFTETAAFVKTPEGEDLKFIPPDFGFEPKHV
ISMIIPQSLEGVKCAPSFLGSAEYGLSFAQIGYAAFGLSMFIKDLGYHAVPIGSDSALSIPIA
IQAGLGEYSRSGQMITPEFGPNVRLCEVFTDMPLNHDKPISFGVTEFCKTCKKCAEACPP
QAISYEDPTIDGPRGQMHNSGIKRWYVDPVKCFEFWSRDNVRNCCGACIAACPFTKPEA
WHHTLIRSLVGAPVITPFMKDMDDIFGYGKPNEKAKADWWK

> TmrA | *Dehalobacter* sp. UNSWDHB codon optimized nucleotide

**ATGGATAAGGAAAAAAGTAATAATGACAAACCTGCCACAAAAATCAATCGAAG
ACAGTTTCTTAAATTCGGCGCAGGCGCGAGTTCTGGCATAGCGATAGCGACAG
CGGCAACTGCGTTAGGAGGTAAAAGTCTA**ATCGACCCTAAGCAAGTCTACGCCGG
CACAGTGAAAGAACTAGACGAATTGCCTTTTAACATCCCGGCGGATTACAAGCCTTT
CACGAATCAGCGCAATATATTCGGACAAGCCGTCTTGGGGGTGCCTGAACCACTTGC
TCTAGTGGAACGATTTGATGAGGTTCGATGGAACGGGTGGCAGACAGATGGCTCTC
CGGGATTGACAGTCTTGGACGGAGCAGCGGCGCGCTAGCTTCGCGGTAGATTATT
ATTTCAACGGTGAAAATTCTGCGTGTAGAGCCAACAAAGGCTTTTTCGAATGGCACC
CGAAAGTCCCTGAGCTTAACTTCAAGTGGGGAGATCCAGAGCGTAATATCCACAGC
CCTGGAGTGAAAAGTGCCGAAGAGGGCACAATGGCTGTTAAGCGAATGGCTCGATT
TTTCGGAGCGGCTAAAGCGGGGATTGCTCCGTTTGATAAGCGATGGGTATTTACGGA
AACGGCAGCATTTGTAAAGACACCAGAGGGAGAAGACTTAAAGTTTATTCCGCCGG
ATTTCGGCTTCGAGCCTAAGCATGTGATAAGTATGATAATTCCGCAGTCATTAGAGG
GCGTTAAATGCGCTCCTTCTTTTTTAGGAAGCGCCGAGTACGGTCTTAGTTTCGCGCA
GATTGGTTACGCTGCTTTTGGGCTTAGCATGTTTATTAAGGACTTAGGTTACCACGCA
GTTCCAATTGGGTCAGACTCTGCCTTATCAATACCTATTGCGATCCAAGCAGGCCTT

GGGGAGTATAGTAGATCAGGGCAGATGATTACTCCGGAGTTTGGACCTAATGTACG
ATTATGTGAAGTTTTCACCGACATGCCTTTGAATCATGACAAGCCTATAAGTTTCGG
GGTGACGGAGTTTTGTAAGACTTGTAAAAAGTGTGCAGAGGCATGCCCACCTCAGG
CGATAAGCTACGAGGACCCTACTATTGACGGACCTAGAGGCCAAATGCATAATAGT
GGAATAAAGCGCTGGTATGTCGATCCTGTGAAGTGTTTTGAATTTTGGAGCCGTGAT
AACGTCCGTAACTGTTGCGGTGCTTGCATCGCCGCATGTCCATTCACCAAACCGGAG
GCTTGGCACCACACTCTAATCCGCAGTTTAGTTGGCGCACCTGTTATCACTCCTTTTA
TGAAAGATATGGATGACATCTTTGGCTATGGCAAACCTAATGAGAAGGCCAAAGCT
GATTGGTGGAAGTAA

> CfrA | *Dehalobacter* sp. CF amino acid

**MDKEKSNNDKPATKINRRQFLKFGAGASSGIAIATAATALGGKSL**IDPKQVYAGTV
KELDELPFNIPADYKPFTNQRNIYGQAVLGVPEPLALVERFDEVRWNGWQTDGSPGLTV
LDGAAARASFAVDYYFNGENSACRANKGFFEWHPKVAELNFKWGDPERNIHSPGVKS
AEEGTMAVKKIARFFGAAKAGIAPFDKRWVFTETYAFVKTPEGESLKFIPPDFGFEPKHV
ISMIIPQSPEGVKCDPSFLGSTEYGLSCAQIGYAAFGLSMFIKDLGYHAVPIGSDSALAIPI
AIQAGLGEYSRSGLMITPEFGSNVRLCEVFTDMPLNHDKPISFGVTEFCKTCKKCAEACA
PQAISYEDPTIDGPRGQMQNSGIKRWYVDPVKCLEFMSRDNVGNCCGACIAACPFTKPE
AWHHTLIRSLVGAPVITPFMKDMDDIFGYGKLNDEKAIADWWK

> CfrA | *Dehalobacter* sp. CF codon optimized nucleotide

**ATGGACAAGGAAAAAAGTAACAACGATAAGCCGGCAACAAAAATTAATCGCAG
ACAATTCCTTAAATTTGGAGCTGGAGCTTCTTCGGGTATTGCAATTGCCACTGC
AGCTACTGCATTGGGAGGGAAATCACTT**ATCGATCCCAAACAGGTATATGCTGGA
ACGGTCAAGGAACTGGATGAACTTCCCTTTAATATCCCGGCAGACTACAAACCGTTT
ACCAATCAAAGGAATATATATGGCCAGGCTGTATTGGGAGTACCCGAACCTCTAGC
ACTTGTAGAGCGTTTTGATGAAATAAGATGGAATGGTTGGCAGACAGATGGTTCGCC
CGGTCTTACTGTACTTGATGGTGCGGCTGCTCGTGCAAGCTTTGCCGTTGATTATTAT
TTTAACGGGGAAAATAGCGCCTGCAGGGCCAATAAAGGTTTTTTTGAATGGCATCCC
AAAGTGGCCGAGCTGAACTTTAAGTGGGGCGATCCGGAGAGAAATATTCATTCCCC
CGGTGTAAAAAGTGCCGAAGAAGGAACGATGGCAGTAAAAAAAATAGCTAGATTTT
TCGGCGCTGCTAAAGCTGGGATAGCGCCTTTTGACAAACGTTGGGTTTTTACTGAAA
CGTATGCCTTTGTTAAAACGCCTGAGGGTGAAAGTCTGAAATTTATCCCTCCGGATT
TTGGGTTTGAGCCCAAGCATGTAATCTCGATGATTATCCCACAGTCGCCAGAAGGAG
TAAAGTGTGACCCGTCCTTTTTAGGATCAACTGAATATGGATTAAGTTGTGCCCAGA
TTGGATATGCTGCATTCGGTTTATCCATGTTTATTAAAGATCTGGGATATCATGCGGT
TCCAATCGGATCTGACAGTGCATTAGCTATACCTATAGCTATTCAGGCGGGTCTGGG
GGAATACAGCAGGTCGGGGCTAATGATTACGCCTGAATTTGGTTCAAATGTTAGACT
CTGTGAAGTATTTACTGACATGCCTTTAAATCATGATAAACCTATTTCATTCGGAGTA
ACTGAATTTTGCAAAACCTGCAAAAAATGCGCTGAAGCATGCGCCCCTCAAGCTATT
AGCTATGAAGATCCTACCATTGATGGACCTCGTGGGCAAATGCAAAATTCGGGAAT
AAAGAGATGGTATGTTGACCCGGTGAAGTGCTTAGAATTCATGTCGCGTGATAACGT
CAGAAACTGCTGCGGAGCTTGTATAGCTGCTTGCCCATTTACTAAGCCGGAAGCCTG
GCACCATACCTTAATTAGGAGTCTAGTAGGAGCACCTGTTATTACTCCATTCATGAA

AGATATGGATGATATTTTTGGATACGGAAAGCCGAATGATGAAAAAGCGATAGCAG
ATTGGTGGAAATAA

> DcrA | *Dehalobacter* sp. DCA amino acid

**MDKEKSNNDKPATKINRRQFLKFGAGASSGIAIATAATALGGKSL**IDPKQVYAGTV
KELDELPFNIPADYKPFTHQRNIWGQALLGVPEPLALRERFAEVRWNGWQTDGSPGLTV
LDGAAAHASWAVDYYLNGENSACRANKGFFEWHPKVPELNFRWGDPERNIHSPGVKS
AEEGTMAVKKIARFFGAAKAGIAPFDKRWVFTETAAFVKTPEGESLKFIPPDFGFEPKHV
ISMIIPQSLEGTKCAPSFLGSAEYGLSYTQIGYAAFGLSMFIKDLGYHAVPIGADSALAIPI
AIQAGLGEYSRSGLMITPEFGPNVRLCEVFTDMPLNHDKPISFGVTEFCKTCKKCAEACA
PQAISYEDPTIDGPRGQMQNSGIKRWYVDPVKCFEFWSRDNVRNCCGACIAACPFTKPE
AWHHTLIRSLVGAPVITPFMKDMDDIFGYGKPNDEKAIADWWK

> DcrA | *Dehalobacter* sp. DCA codon optimized nucleotide

**ATGGACAAGGAAAAAAGTAACAACGATAAGCCGGCAACAAAAATTAATCGCAG
ACAATTCCTTAAATTTGGAGCTGGAGCTTCTTCGGGTATTGCAATTGCCACTGC
AGCTACTGCATTGGGAGGGAAATCACTT**ATCGATCCCAAACAGGTATATGCTGGA
ACGGTCAAGGAACTGGATGAACTTCCCTTTAATATCCCGGCAGACTACAAACCGTTT
ACCCATCAAAGGAATATATGGGGCCAGGCTTTATTGGGAGTACCCGAACCTCTAGC
ACTCAGAGAGCGTTTTGCTGAAGTAAGATGGAATGGTTGGCAGACAGATGGTTCGC
CCGGTCTTACTGTACTTGATGGTGCGGCTGCTCATGCAAGCTGGGCCGTTGATTATT
ATCTTAACGGGGAAAATAGCGCCTGCAGGGCCAATAAAGGTTTTTTGAATGGCATC
CCAAAGTGCCCGAGCTGAACTTTAGGTGGGGCGATCCGGAGAGAAATATTCATTCC
CCCGGTGTAAAAAGTGCCGAAGAAGGAACGATGGCAGTAAAAAAAATAGCTAGATT
TTTCGGCGCTGCTAAAGCTGGGATAGCGCCTTTTGACAAACGTTGGGTTTTTACTGA
AACGGCTGCCTTTGTTAAAACGCCTGAGGGTGAAAGTCTGAAATTTATCCCTCCGGA
TTTTGGGTTTGAGCCCAAGCATGTAATCTCGATGATTATCCCACAGTCGCTAGAAGG
AACAAAGTGTGCCCCGTCCTTTTTAGGATCAGCTGAATATGGATTAAGTTATACCCA
GATTGGATATGCTGCATTCGGTTTATCCATGTTTATTAAAGATCTGGGATATCATGCG
GTTCCAATCGGAGCTGACAGTGCATTAGCTATACCTATAGCTATTCAGGCGGGTCTG
GGGGAATACAGCAGGTCGGGGCTAATGATTACGCCTGAATTTGGTCCAAATGTTAG
ACTCTGTGAAGTATTTACTGACATGCCTTTAAATCATGATAAACCTATTTCATTCGGA
GTAACTGAATTTTGCAAAACCTGCAAAAAATGCGCTGAAGCATGCGCCCCTCAAGCT
ATTAGCTATGAAGATCCTACCATTGATGGACCTCGTGGGCAAATGCAAAATTCGGGA
ATAAAGAGATGGTATGTTGACCCGGTGAAGTGCTTTGAATTCTGGTCGCGTGATAAC
GTCAGAAACTGCTGCGGAGCTTGTATAGCTGCTTGCCCATTTACTAAGCCGGAAGCC
TGGCACCATACCTTAATTAGGAGTCTAGTAGGAGCACCTGTTATTACTCCATTCATG
AAAGATATGGATGATATTTTTGGATACGGAAAGCCGAATGATGAAAAAGCGATAGC
AGATTGGTGGAAATAA

> HchA | *Dehalobacter* sp. HCH1 amino acid

**<u>MISENDKEKKTQQKKSKEVSRRGFLKASLGVGVGATGAALFGSELPGSNIIANA</u>**AT
VEHDTMPVEISADYKRYSSASLLPLNQGVKELYEVRAGVVPPPEGWGWDPVNTALFHA
AWSIEGDINHYKPSAGYRSTEGLCTWDGKVNPDTHKFESPEAASDAVKRAAMFFGASK
VGIAPYDERWLHSEIMDDPATNNLIPNDLPFTPTHAIVILLEMDYDGSEAAPLPPATIATA
DIISNLGIINHKMATFIRQLGYKCLPCTNDTALSIPMAIQAGLGEMSRAGILLTSEFGARV
MICKLFVDMPLAADKPISFGGTEFCKTCMKCADACPTQAISHDKEPSYKVFAATNPGVK
KWAMDGTKCITQLATTGGYCSICIKVCPYNKQQEWHHEFVKLGTKTPARPVLRFFDDL
FGYGKITPEDAVKKFWKK

> HchA | *Dehalobacter* sp. HCH1 codon optimized nucleotide, TAT removed and start codon added

ATGGCCACGGTTGAGCATGACACTATGCCAGTCGAGATCTCGGCGGACTACAAACG
CTATTCCAGCGCTTCGCTGTTGCCTTTAAACCAGGGCGTGAAAGAGCTGTACGAGGT
GCGTGCGGGTGTTGTTCCACCGCCGGAAGGCTGGGGTTGGGATCCGGTTAATACTGC
TCTGTTTCATGCCGCTTGGAGTATCGAGGGCGATATCAATCACTATAAGCCTAGCGC
AGGATACCGCTCAACGGAAGGGCTGTGTACTTGGGACGGAAAGTCAACCCGGATA
CGCATAAATTCGAATCTCCAGAAGCAGCAAGTGATGCGGTTAAACGTGCGGCCATG
TTTTTTGGTGCTAGCAAGGTGGGCATTGCACCGTACGATGAACGTTGGCTGCATTCG
GAAATTATGGACGACCCGGCTACAAATAATCTGATTCCGAACGATTTACCGTTTACG
CCGACGCATGCTATTGTGATTTTGCTCGAAATGGATTACGATGGAAGCGAAGCTGCC
CCTTTGCCGCCGGCAACGATTGCGACAGCAGATATTATTAGCAATCTGGGCATCATC
AACCATAAAATGGCCACTTTTATTCGCCAACTGGGCTACAAGTGCCTGCCGTGTACT
AACGACACTGCACTGAGTATCCCAATGGCAATTCAAGCTGGGCTGGGCGAGATGAG
CCGTGCTGGTATTCTGCTGACCTCCGAATTTGGTGCCCGTGTAATGATCTGTAAACTG
TTTGTTGACATGCCCTTAGCCGCCGATAAACCAATCAGCTTTGGCGGCACTGAATTC
TGTAAAACGTGTATGAAATGCGCTGATGCGTGTCCTACCCAGGCGATTTCACATGAT
AAAGAGCCGAGCTATAAAGTATTTGCGGCGACAAACCCAGGGGTTAAGAAGTGGGC
CATGGACGGAACCAAATGCATCACTCAGCTGGCGACCACGGGCGGTTACTGCTCGA
TCTGTATTAAGGTCTGTCCTTACAATAAGCAGCAGGAATGGCATCATGAATTCGTCA
AACTGGGAACTAAAACGCCGGCCCGTCCGGTACTGCGTTTTTTTGATGACCTGTTCG
GTTACGGCAAAATTACCCCAGAGGATGCGGTTAAAAAATTTTGGAAAAAA

> DHB14 | *Dehalobacter* sp. CF amino acid

<u>MGEINRRNFLKASMLGAAAAAVASASVVKGMVSPLVADA</u>ADIVAPITETSEFPYKVDA
KYQRYNCMKNFFEKTFDPEENKTPIKFHNDDVSKITGKKDTGKDLPTLNAERLGIKGRP
ATHTETGVLFYSQHMGVMPPQRSKETGWTSLDEALNAGAWAVEFDFSGFNATVGGGP
GSLIPSYPINPMTNEMANDPVLVSGLYSWDNSDAEGVRQQNQQWKFKSKEEASKIVKK
AACFLGADLVGIAPYDDRWTYASWGRDIEKPFKLPNGKIKYLPWDLPKMLSGGGIEVF
GHTEFESDWEKYGGFKPKSVIVFVFEMDIEALRTSPSVIASAAAGKAYSSMGEVSYKIAV
FLRKLGYYATSSGNDTGLNVPLAVQAGLGEAGRNGLLITQKFGPRHRIAKVYTDLELAP
DKPRKFGVREFCRLCKKCADACPAQAISHEKDPKVLQPGDCEESENPYTEKWHVDGER

CGSFWTYNGSPCANCVAVCSWNKLETWNHDVARIATQIPLIQDAARKFDEWFGYNGP
VNPDERLESGYVQNMVKDFWNNPESIK

> DHB14 | *Dehalobacter* sp. CF nucleotide

ATGGGCGAAATCAACAGGAGGAATTTTTTAAAAGCCTCGATGCTTGGAGCAGCTGC
AGCTGCTGTAGCCTCGGCATCTGTGGTGAAGGGGATGGTTAGCCCCTTGGTGGCTGA
TGCTGCGGACATCGTGGCTCCGATCACGGAAACCTCAGAATTTCCATACAAGGTGGA
TGCAAAGTACCAGCGTTACAATTGTATGAAGAACTTCTTTGAAAAGACTTTCGATCC
GGAAGAAAACAAGACTCCTATTAAGTTTCATAACGACGATGTTTCCAAAATCACAG
GCAAGAAAGATACGGGGAAAGACCTGCCCACGCTTAATGCGGAAAGACTTGGGATC
AAAGGGCGTCCGGCGACACATACAGAAACAGGCGTTCTTTTCTATAGTCAGCATATG
GGTGTCATGCCACCCCAGCGCAGTAAAGAAACAGGCTGGACCTCGTTGGATGAGGC
CTTGAACGCCGGAGCCTGGGCTGTAGAATTTGATTTTTCCGGATTTAACGCAACAGT
TGGTGGTGGTCCGGGAAGTTTAATCCCGTCTTATCCCATAAATCCTATGACCAATGA
AATGGCTAATGATCCTGTCCTGGTATCCGGGTTGTACAGCTGGGACAATAGCGATGC
TGAAGGTGTGCGACAACAAAACCAACAGTGGAAATTCAAATCAAAGGAAGAAGCC
AGTAAAATCGTTAAAAAAGCAGCCTGTTTCCTGGGAGCCGATCTGGTCGGCATCGCA
CCCTATGACGATCGTTGGACCTATGCTTCCTGGGGCAGGGATATTGAAAAACCTTTT
AAACTACCCAACGGCAAAATCAAATATCTGCCATGGGATTTGCCAAAGATGCTGTC
AGGCGGGGGAATAGAAGTTTTCGGACATACAGAATTTGAATCTGATTGGGAGAAGT
ATGGAGGTTTCAAACCAAAAAGTGTGATTGTCTTCGTTTTTGAAATGGATATTGAAG
CTTTGCGCACCTCACCGTCAGTCATTGCCAGTGCCGCGGCGGGGAAAGCTTACTCCA
GTATGGGAGAAGTATCTTACAAAATCGCGGTTTTTCTGAGGAAGCTTGGCTATTATG
CAACGTCGTCAGGAAACGATACCGGATTGAATGTTCCTTTGGCCGTTCAGGCCGGGC
TTGGAGAAGCAGGCAGAAACGGACTTTTAATTACCCAGAAATTCGGTCCGAGACAT
CGTATCGCCAAAGTCTACACCGACCTGGAACTTGCTCCGGACAAGCCGAGAAAATT
CGGGGTACGCGAGTTCTGCCGCCTGTGCAAAAAATGTGCGGATGCCTGCCCCGCCC
AGGCCATCTCCCATGAGAAAGACCCTAAGGTTCTGCAGCCAGGGGATTGTGAGGAA
TCCGAAAATCCATATACTGAAAAATGGCATGTTGATGGCGAACGCTGCGGCTCTTTC
TGGACCTATAACGGTAGTCCCTGCGCAAATTGTGTAGCTGTATGTTCGTGGAACAAA
CTCGAGACCTGGAACCACGACGTGGCCAGAATAGCCACCCAAATACCATTGATTCA
GGATGCAGCCCGCAAGTTTGATGAGTGGTTCGGCTATAACGGGCCTGTAAACCCTGA
TGAAAGACTTGAATCGGGTTATGTTCAGAACATGGTAAAAGACTTCTGGAATAATCC
TGAGTCTATAAAATAA

> DHB15 | *Dehalobacter* sp. DCA amino acid

MGEINRRNFLKASMLGAAAAAVASASAVKGMVSPLVADAADIVAPITETSEFPYKVDA
KYQRYNSLKNFFEKAFDPEANKTPIKFHYDDVSKITGKKDTGKDLPTLNAERLGIKGRP
ATHTETAMLFFTQHFGAMPPQRHNEAGWTPVEAALNAGAWAVEFDFSGFNATGGGPG
SLIPSYPINPMTNEMAKEPVIVSGLYNWDNSDAEGVRQQGQQWKFKSKEEASKIVKKSV
KFLGADLVGIAPYDERWTYSNWGREIPKPFKMPDGRTKYFPWDLPKMMSGGGVEVFG
HAEFEPDWEKYGGFKPKSVIVFILEEDYEAIRTSPSVIASATVGKTYSNMGEVAYKIAVF
LRKLGYYAVPAGNDTGMSVPMAVQAGLGEAGRNGLLITQKFGPRHRIAKVYTDLELAP
DKPKKFGVREFCRLCKKCADACPAQAISHEKDPKVLQPEDVEVSENPYTEKWYVDSER

CGSFWAYNGSPCVNCVAVCSWNKVETWNHDVARIATRIPLLQDAARKFDEWFGYNGP
VNPDERLESGYVQNMVKDFWNNPESIKQ

> DHB15 | *Dehalobacter* sp. DCA nucleotide

ATGGGAGAAATCAACAGGAGGAATTTTTTAAAAGCCTCGATGCTTGGAGCAGCTGC
AGCTGCTGTAGCCTCGGCATCTGCGGTGAAGGGGATGGTTAGCCCCTTGGTGGCTGA
TGCTGCGGACATCGTGGCTCCGATCACGGAAACCTCAGAATTTCCATACAAGGTGGA
TGCAAAGTACCAACGTTACAATAGTCTGAAGAACTTCTTTGAAAAGGCTTTCGACCC
GGAAGCAAACAAGACTCCTATTAAGTTTCATTACGACGATGTTTCCAAAATCACAGG
CAAGAAAGATACGGGGAAAGACCTGCCCACGCTTAATGCGGAAAGACTTGGGATCA
AAGGGCGTCCGGCGACACATACAGAAACAGCCATGCTTTTCTTTACTCAACATTTTG
GTGCCATGCCACCCCAGCGCCATAATGAAGCAGGCTGGACCCCAGTGGAAGCGGCC
TTGAACGCCGGAGCCTGGGCTGTAGAATTTGATTTTTCCGGATTTAACGCAACAGGT
GGTGGTCCGGGAAGTTTAATCCCGTCTTATCCCATAAATCCTATGACCAATGAAATG
GCTAAAGAGCCTGTCATAGTATCCGGTTTGTACAACTGGGACAATAGCGATGCTGAA
GGTGTGCGACAACAAGGCCAACAGTGGAAATTCAAATCAAAGGAAGAAGCCAGTA
AAATCGTTAAAAAGTCAGTCAAGTTCCTGGGAGCCGATCTGGTCGGCATCGCACCCT
ATGACGAGCGTTGGACCTATTCTAACTGGGGCAGGGAGATTCCAAAACCTTTTAAAA
TGCCCGACGGCAGAACTAAATATTTTCCATGGGATTTGCCAAAGATGATGTCAGGCG
GGGGAGTAGAAGTTTTCGGACATGCAGAATTTGAACCTGATTGGGAGAAGTATGGA
GGTTTCAAACCAAAAAGTGTGATTGTCTTCATTCTTGAAGAGGATTACGAAGCTATA
CGCACCTCACCGTCAGTCATTGCCAGTGCCACGGTGGGGAAAACTTACTCCAATATG
GGAGAAGTAGCTTACAAAATCGCTGTTTTTCTGAGGAAGCTTGGCTATTATGCAGTG
CCGGCAGGAAACGATACCGGAATGAGTGTTCCTATGGCCGTTCAGGCCGGGCTTGG
AGAAGCAGGTAGAAACGGACTTTTAATTACCCAGAAATTCGGTCCGAGACATCGCA
TCGCCAAAGTCTACACCGACCTGGAACTTGCTCCGGACAAGCCGAAAAAATTCGGG
GTACGCGAGTTCTGCCGCCTGTGCAAAAAATGTGCGGATGCCTGCCCCGCCCAGGCC
ATCTCCCACGAGAAAGACCCTAAGGTTCTGCAGCCAGAGGATGTTGAGGTATCCGA
AAATCCATATACTGAAAAATGGTATGTTGATTCCGAACGCTGCGGCTCTTTCTGGGC
CTATAACGGTAGTCCCTGCGTAAATTGTGTAGCTGTATGTTCGTGGAACAAAGTCGA
GACCTGGAACCACGATGTGGCCAGAATAGCCACTCGAATACCATTGCTTCAGGATG
CAGCCCGCAAGTTTGATGAATGGTTCGGCTATAACGGGCCTGTAAACCCTGATGAAA
GACTTGAATCGGGTTATGTTCAGAACATGGTCAAAGACTTCTGGAATAATCCTGAGT
CTATAAAACAATAA

> VcrA | *Dehalococcoides* sp. VC amino acid

**MSKFHKTISRRDFMKGLGLAGAGIGAVAASAPVFHDIDELVSSE**ANSTKDQPWYVK
HREHFDPTITVDWDIFDRYDGYQHKGVYEGPPDAPFTSWGNRLQVRMSGEEQKKRILA
AKKERFPGWDGGLHGRGDQRADALFYAVTQPFPGSGEEGHGLFQPYPDQPGKFYARW
GLYGPPHDSAPPDGSVPKWEGTPEDNFLMLRAAAKYFGAGGVGALNLADPKCKKLIYK
KAQPMTLGKGTYSEIGGPGMIDAKIYPKVPDHAVPINFKEADYSYYNDAEWVIPTKCESI
FTFTLPQPQELNKRTGGIAGAGSYTVYKDFARVGTLVQMFIKYLGYHALYWPIGWGPG
GCFTTFDGQGEQGRTGAAIHWKFGSSQRGSERVITDLPIAPTPPIDAGMFEFCKTCYICRD
VCVSGGVHQEDEPTWDSGNWWNVQGYLGYRTDWSGCHNQCGMCQSSCPFTYLGLEN
ASLVHKIVKGVVANTTVFNSFFTNMEKALGYGDLTMENSNWWKEEGPIYGFDPGT

> VcrA | *Dehalococcoides* sp. VC codon optimized nucleotide

**ATGAGTAAATTTCATAAAACGATTAGCCGCCGAGATTTCATGAAAGGACTAGGA
TTAGCCGGGGCAGGCATAGGCGCTGTTGCGGCGTCAGCTCCGGTTTTTCATGA
CATTGATGAACTTGTTTCAAGCGAAG**CAAATTCTACTAAAGATCAACCTTGGTAC
GTTAAGCATCGAGAGCATTTTGACCCTACGATTACAGTTGACTGGGATATTTTTGAT
AGATATGACGGGTATCAGCATAAGGGTGTCTATGAAGGCCCTCCAGATGCTCCCTTT
ACATCATGGGGCAATAGGCTTCAGGTGAGAATGTCAGGTGAAGAGCAAAAGAAGCG
AATTTTGGCCGCTAAAAAAGAGAGGTTCCCTGGTTGGGACGGTGGGTTACACGGGA
GAGGGGATCAGCGGGCGGATGCACTATTTTACGCAGTAACTCAACCATTTCCTGGTA
GTGGTGAGGAAGGGCACGGACTATTCCAACCTTATCCTGATCAACCCGGTAAGTTTT
ACGCGAGATGGGGTTTGTATGGTCCGCCACATGATTCAGCGCCACCTGATGGGAGC
GTACCAAAATGGGAGGGTACTCCAGAAGACAATTTTCTAATGCTGAGGGCAGCTGC
AAAATATTTTGGTGCTGGTGGCGTTGGTGCTCTTAACCTGGCAGATCCCAAATGCAA
AAAACTAATATATAAGAAAGCTCAGCCGATGACTCTAGGAAAAGGAACATACAGTG
AAATAGGTGGACCAGGAATGATCGATGCAAAAATTTATCCCAAGGTTCCTGACCAT
GCCGTACCTATTAACTTTAAGGAAGCGGATTATAGCTACTACAATGATGCAGAGTGG
GTTATTCCAACAAAGTGTGAATCCATTTTCACTTTCACCCTACCTCAACCACAAGAA
CTCAATAAGAGGACGGGTGGTATAGCAGGTGCTGGATCATATACTGTATACAAAGA
TTTCGCTAGGGTAGGCACTTTAGTCCAAATGTTTATTAAGTATCTAGGTTATCACGCT
TTATATTGGCCAATTGGATGGGGACCGGGTGGTTGCTTTACCACTTTTGACGGGCAA
GGTGAACAGGGTAGAACAGGTGCTGCTATCCATTGGAAGTTTGGTTCTTCACAACGT
GGTTCTGAAAGAGTAATAACTGATTTACCGATAGCTCCTACCCCGCCAATTGATGCA
GGTATGTTTGAGTTTTGCAAAACCTGTTATATATGCCGTGACGTTTGCGTCTCTGGGG
GTGTGCACCAAGAAGACGAACCAACTTGGGATTCAGGTAATTGGTGGAATGTACAA
GGATATCTCGGCTACCGAACGGATTGGAGTGGTTGCCATAACCAGTGCGGTATGTGT
CAATCCTCCTGCCCTTTTACTTATTTAGGTTTGGAAAATGCTTCATTAGTGCACAAAA
TAGTAAAAGGTGTTGTTGCTAACACGACTGTTTTTAATAGTTTTTTTACCAATATGGA
GAAAGCATTAGGATATGGTGATTTAACCATGGAAAATTCTAACTGGTGGAAAGAAG
AAGGACCGATATACGGCTTTGATCCCGGTACTTAG

> BvcA | *Dehalococcoides* sp. BAV1 amino acid

**MHNFHCTISRRDFMKGLGLAGAGIGAATSVMPNFHDLDEVISAASA**ETSSLSGKSLN
NFPWYVKERDFENPTIDIDWSILARNDGYNHQGAYWGPVPENGDDKRYPDPADQCLTL
PEKRDLYLAWAKQQFPDWEPGINGHGPTRDEALWFASSTGGIGRYRIPGTQQMMSTMR
LDGSTGGWGYFNQPPAAVWGGKYPRWEGTPEENTLMMRTVCQFFGYSSIGVMPITSNT
KKLFFEKQIPFQFMAGDPGVFGGTGNVQFDVPLPKTPVPIVWEEVDKGYYNDQKIVIPN
KANWVLTMTMPLPEDRFKRSLGWSLDASSMIAYPQMAFNGGRVQTFLKALGYQGLGG
DVAMWGPGGAFGVMSGLSEQGRAANEISPKYGSATKGSNRLVCDLPMVPTKPIDAGIH
KFCETCGICTTVCPSNAIQVGPPQWSNNRWDNTPGYLGYRLNWGRCVLCTNCETYCPF
FNMTNGSLIHNVVRSTVAATPVFNSFFRQMEHTFGYGMKDDLNDWWNQSHKPW

> BvcA | *Dehalococcoides* sp. BAV1 codon optimized nucleotide

**ATGCATAATTTCCATTGTACGATAAGTAGGCGAGATTTTATGAAGGGATTGGGG
TTAGCGGGAGCAGGGATAGGTGCCGCGACTTCAGTTATGCCGAATTTTCACGA
CTTGGATGAAGTAATTTCTGCTGCTAGTGCC**GAAACCAGTTCTTTGTCGGGTAAA
TCTCTTAATAATTTTCCTTGGTATGTGAAAGAAAGGGATTTTGAAAATCCTACCATTG

ATATAGATTGGTCTATACTTGCGCGTAATGACGGTTACAATCATCAGGGAGCCTATT
GGGGACCTGTACCTGAAAATGGAGATGATAAAAGGTATCCTGATCCCGCGGACCAG
TGTCTTACTCTACCAGAAAAGAGAGATCTTTATTTAGCGTGGGCAAAACAGCAATTT
CCTGACTGGGAACCAGGAATTAATGGCCATGGGCCAACAAGGGACGAAGCTTTATG
GTTTGCCTCAAGTACAGGTGGTATCGGTAGGTATAGAATTCCTGGTACCCAGCAAAT
GATGTCCACAATGCGTCTTGACGGGTCTACTGGTGGTTGGGGTTATTTCAATCAACC
ACCGGCAGCAGTCTGGGGAGGGAAATACCCAAGGTGGGAAGGAACTCCTGAAGAA
AATACGTTGATGATGCGAACTGTTTGTCAATTTTTTGGTTACTCCAGTATAGGTGTAA
TGCCAATCACCAGCAATACAAAGAAGCTTTTTTTTGAAAAGCAAATACCTTTCCAAT
TTATGGCTGGAGATCCCGGTGTATTTGGGGGAACGGAAATGTGCAGTTTGATGTCC
CGCTGCCAAAGACACCTGTTCCAATAGTCTGGGAGGAAGTCGATAAAGGGTATTAT
AATGACCAGAAAATTGTAATACCCAATAAGGCTAACTGGGTATTAACAATGACAAT
GCCTTTACCAGAAGATCGTTTTAAACGTTCTCTAGGGTGGTCACTTGACGCTTCAAG
TATGATTGCCTATCCTCAGATGGCTTTTAATGGAGGCCGAGTTCAGACTTTTTTAAAA
GCACTTGGCTATCAAGGACTTGGTGGCGACGTGGCTATGTGGGGACCTGGTGGTGCT
TTTGGAGTTATGAGTGGTCTTTCCGAACAAGGTCGTGCTGCTAATGAAATCAGCCCC
AAATACGGTTCGGCAACTAAGGGCTCTAATCGATTAGTTTGTGATTTGCCCATGGTT
CCGACCAAGCCAATTGATGCTGGCATACACAAATTCTGTGAAACGTGTGGCATTTGT
ACAACAGTTTGTCCCTCAAATGCTATCCAGGTAGGTCCTCCACAATGGAGTAATAAT
CGGTGGGATAATACCCCTGGTTATCTTGGTTATCGACTTAACTGGGGTAGATGTGTT
CTTTGTACAAACTGTGAGACCTATTGCCCATTTTTTAACATGACTAATGGTTCTTTGA
TTCATAACGTAGTCAGATCCACAGTTGCAGCTACACCGGTTTTTAATTCATTTTTCCG
CCAAATGGAACATACATTTGGATATGGTATGAAAGATGATTTAAACGATTGGTGGA
ATCAATCACACAAGCCTTGGTAA

> TceA | *Dehalococcoides* sp. 195 amino acid

**MSEKYHSTVTRRDFMKRLGLAGAGAGALGAAVLA**ENNLPHEFKDVDDLLSAGKAL
EGDHANKVNNHPWWVTTRDHEDPTCNIDWSLIKRYSGWNNQGAYFLPEDYLSPTYTG
RRHTIVDSKLEIELQGKKYRDSAFIKSGIDWMKENIDPDYDPGELGYGDRREDALIYAAT
NGSHNCWENPLYGRYEGSRPYLSMRTMNGINGLHEFGHADIKTTNYPKWEGTPEENLLI
MRTAARYFGASSVGAIKITDNVKKIFYAKVQPFCLGPWYTITNMAEYIEYPVPVDNYAIP
IVFEDIPADQGHYSYKRFGGDDKIAVPNALDNIFTYTIMLPEKRFKYAHSIPMDPCSCIAY
PLFTEVEARIQQFIAGLGYNSMGGGVEAWGPGSAFGNLSGLGEQSRVSSIIEPRYGSNTK
GSLRMLTDLPLAPTKPIDAGIREFCKTCGICAEHCPTQAISHEGPRYDSPHWDCVSGYEG
WHLDYHKCINCTICEAVCPFFTMSNNSWVHNLVKSTVATTPVFNGFFKNMEGAFGYGP
RYSPSRDEWWASENPIRGASVDIF

> TceA | *Dehalococcoides* sp. 195 codon optimized nucleotide, TAT removed and start codon

ATGGAGAACAACCTGCCGCACGAATTCAAAGACGTTGACGATCTGCTGTCCGCCGG
TAAGGCGCTGGAAGGAGACCATGCGAACAAGGTGAATAACCACCCGTGGTGGGTTA
CCACCCGTGATCACGAGGACCCAACGTGTAACATCGATTGGAGCCTCATTAAACGCT
ATTCCGGTTGGAATAATCAAGGCGCGTACTTTCTCCCGGAAGATTACTTATCCCCGA
CTTACACTGGCCGCCGTCACACCATCGTAGACTCTAAATTAGAAATTGAACTGCAAG
GAAAAAAATACCGTGATAGCGCTTTCATCAAGTCTGGTATTGACTGGATGAAAGAA
AACATCGATCCGGATTACGATCCGGGGGAACTGGGGTACGGCGATCGCCGCGAGGA
CGCGCTGATCTATGCAGCCACCAACGGAAGCCACAACTGCTGGGAAAATCCTTTAT

ATGGCCGCTACGAAGGTAGCCGTCCCTACCTGAGTATGCGCACCATGAATGGAATC
AACGGTCTGCATGAATTCGGGCATGCCGACATCAAAACCACGAATTATCCAAAATG
GGAGGGAACGCCCGAAGAGAATTTACTCATCATGCGGACCGCTGCGCGCTATTTCG
GCGCCAGTTCTGTGGGTGCGATCAAAATCACGGACAACGTGAAGAAAATTTTCTATG
CTAAAGTGCAGCCGTTCTGCTTAGGCCCATGGTATACCATCACGAATATGGCCGAAT
ACATTGAATATCCGGTCCCTGTTGATAACTACGCCATCCCGATTGTATTTGAAGATA
TTCCTGCGGATCAAGGTCATTATTCTTATAAACGCTTTGGTGGCGATGATAAAATTG
CCGTTCCGAACGCACTGGATAATATCTTCACCTACACGATCATGCTTCCGGAAAAAC
GTTTCAAATACGCGCATAGCATCCCGATGGATCCGTGTAGCTGCATCGCATACCCCC
TGTTTACTGAGGTCGAAGCCCGTATTCAGCAATTTATTGCCGGACTGGGTTATAACA
GTATGGGTGGCGGGGTTGAGGCGTGGGGCCCTGGCTCCGCGTTTGGCAATCTGAGC
GGCCTGGGCGAGCAGTCTCGTGTGAGCAGTATTATTGAGCCGCGCTACGGCTCAAAT
ACCAAAGGGTCTCTGCGTATGCTGACCGATCTGCCACTGGCGCCAACCAAACCGATT
GATGCCGGCATCCGTGAGTTCTGTAAAACATGTGGTATTTGTGCAGAACACTGCCCG
ACCCAGGCAATTAGTCACGAAGGTCCGCGTTATGACAGTCCGCATTGGGACTGCGTC
AGTGGTTACGAAGGCTGGCACCTGGACTATCATAAATGTATCAATTGCACCATCTGC
GAGGCGGTGTGCCCATTCTTTACCATGTCAAATAACAGTTGGGTCCACAACCTTGTG
AAATCGACCGTAGCGACAACCCCAGTGTTCAATGGTTTTTTTAAGAACATGGAAGGT
GCGTTTGGATACGGTCCTCGGTACTCCCCTAGCCGTGATGAGTGGTGGGCGTCAGAG
AATCCGATCCGTGGCGCAAGTGTGGACATCTTC

#### Primers, Constructs, and *E. coli* Strains Used

This section includes three tables of the primers, constructs, and *E. coli* strains used in this study.

**Table S3. Primers used for amplification of *rdhA* genes for cloning.** Bolded sequences are complimentary to the *rdhA* gene, non-bolded is an extension for Gibson assembly, and red is the added His tag.

| Primer Name | Sequence (5'→3') | Purpose |
|---|---|---|
| *TmrA_TAT_F* | TTGTATTTCCAGGGC**ATGGACA AGGAAAAAAGTAAC** | Forward primer to amplify TmrA with TAT signal peptide sequence for Gibson Assembly into *p15TV-L* |
| *TmrA_CHis_R* | CAAGCTTCGTCATCAATGGTGAT GGTGGTGATGGCCGCTGCT**TTTC CACCAATCTGCTTT** | Reverse primer to amplify TmrA with an added C-terminal x6His tag for Gibson Assembly into *p15TV-L*. |
| *OG97_no_TAT_F* | TTGTATTTCCAGGGC**ATGATCG ATCCCAAACAGGTA** | Amplify enzymes from OG 97 (TmrA, CfrA, DcrA, etc.), without TAT signal peptide sequence for Gibson Assembly into *p15TVL* |
| *TmrA_R* | CAAGCTTCGTCATCA**TTATTTCC ACCAATCTGCTTTC** | Reverse primer to amplify TmrA for Gibson Assembly into *p15TV-L*. |
| *CfrA_DcrA_R* | CAAGCTTCGTCATCA**TTATTTCC ACCAATCTGCTATC** | Reverse primer to amplify CfrA and DcrA for Gibson Assembly into *p15TV-L*. |
| *VcrA_SUMO_F* | CAATATTGGAAGTGGAGGGATG **GCCAATAGCACAAAAGATCAG** | Forward primer to amplify VcrA without TAT signal peptide sequence for Gibson assembly into *pET-SUMO*. |
| *VcrA_SUMO_R* | AGCCAACTCAGCTTCCTT**TTAGG TGCCCGGATCAAAACC** | Reverse primer to amplify VcrA for Gibson assembly into *pET-SUMO*. |
| *BvcA_SUMO_F* | CAATCCAATATTGGAAGTGGAG GGATG**GAAACAAGTTCACTGTC TGG** | Forward primer to amplify BvcA without TAT signal peptide sequence for Gibson assembly into *pET-SUMO*. |
| *BvcA_SUMO_R* | AGCCAACTCAGCTTCCTT**TTACC AAGGCTTGTGTGACTGG** | Reverse primer to amplify BvcA for Gibson assembly into *pET-SUMO*. |
| *TceA_SUMO_F* | CAATATTGGAAGTGGAGGGATG **GAGAACAACCTGCCGC** | Forward primer to amplify TceA without TAT signal peptide sequence for Gibson assembly into *pET-SUMO*. |
| *TceA_SUMO_R* | AGCCAACTCAGCTTCCTT**TTAGA AGATGTCCACACTTGCGCC** | Reverse primer to amplify TceA for Gibson assembly into *pET-SUMO*. |
| *pET-SUMO_bb_F* | AAGGAAGCTGAGTTGGCT | Forward primer to amplify the backbone of *pET-SUMO*. |
| *pET-SUMO_bb_R* | CCCTCCACTTCCAATATTGGATT G | Reverse primer to amplify the backbone of *pET-SUMO*. |

**Table S4. Expression plasmids used in this work.**

| Plasmid | Encoded Enzyme | Tags | Induction | Source |
|---|---|---|---|---|
| *p15TVL-tmrA-TAT* | TmrA (with TAT sequence) | N-term 6xHis, C-term 6xHis | IPTG | This work |
| *p15TVL-tmrA* | TmrA (no TAT) | N-term 6xHis | IPTG | This work |
| *p15TVL-cfrA* | CfrA (no TAT) | N-term 6xHis | IPTG | This work |
| *p15TVL-dcrA* | DcrA (no TAT) | N-term 6xHis | IPTG | This work |
| *pET21-hchA* | HchA (no TAT) | C-term 6xHis | IPTG | This work |
| *p15TVL-DHB14* | DHB14 (with TAT) | N-term 6xHis | IPTG | Yakunin and Savchenko labs (University of Toronto) |
| *p15TVL-DHB15* | DHB15 (with TAT) | N-term 6xHis | IPTG | Yakunin and Savchenko labs (University of Toronto) |
| *pBAD42-BtuCEDFB* | BtuB, BtuCD, BtuE, BtuF | N/A | Arabinose | Booker lab (Pennsylvania State University) (46) |
| *pET21-vcrA* | VcrA (no TAT) | C-term 6xHis | IPTG | This work |
| *pET21-bvcA* | BvcA (no TAT) | C-term 6xHis | IPTG | This work |
| *pET21-tceA* | TceA (no TAT) | C-term 6xHis | IPTG | This work |
| *p15TVL-vcrA* | VcrA (with TAT) | N-term 6xHis | IPTG | Yakunin and Savchenko labs (University of Toronto) |
| *p15TVL-bvcA* | BvcA (with TAT) | N-term 6xHis | IPTG | Yakunin and Savchenko labs (University of Toronto) |
| *pET-SUMO-vcrA* | SUMO-VcrA (no TAT) | N-term SUMO and x6His | IPTG | This work |
| *pET-SUMO-bvcA* | SUMO-BvcA (no TAT) | N-term SUMO and x6His | IPTG | This work |
| *pET-SUMO-tceA* | SUMO-TceA (no TAT) | N-term SUMO and x6His | IPTG | This work |
| *pG-KJE8* | DnaK-DnaJ-GrpE, GroES-GroEL | N/A | Arabinose, tetracycline | (47) |
| *pG-Tf2* | GroES-GroEL-tig | N/A | Tetracycline | (47) |

**Table S5. *Escherichia coli* strains used in this study.**

| Strain | Description | Source |
|---|---|---|
| *E. coli* DH5α | Cloning strain of *E. coli*. | New England Biolabs |
| *E. coli* BL21(DE3) Lobstr | Expression strain of *E. coli*. | (48) |
| *E. coli* BL21(DE3) Δ*iscR* | Expression strain of *E. coli* with a deletion of the *isc* operon repressor, *iscR*, for increased iron-sulfur cluster production. $Kan^R$ | Hallenbeck lab (University of Montreal) (49, 50) |
| *E. coli* SufFeScient (BL21λDE3 *cat*-$P_{sufA}$(Fur*)) | Expression strain of *E. coli* with a repaired and overexpressed *suf* operon, for increased iron-sulfur cluster production. $Cm^R$ | Antony lab (St. Louis University School of Medicine) and Kiley lab (University of Wisconsin-Madison) (51) |
| *E. coli ara-Suf* Δ*iscR* (BL21λDE3 *cat-araC*-$P_{BAD}$-*suf*, Δ*iscR*::*kan*, Δ*himA*::$Tet^R$) | Expression strain of *E. coli* with a deletion of the *isc* operon repressor, *iscR*, and *suf* expression under arabinose control for increased iron-sulfur cluster production. $Cm^R$, $Kan^R$, $Tet^R$ | |

#### SI Text 3 – Expression, Purification, and Assay Data

This section includes figures and tables with additional information about the expression of TmrA, the purification of both TmrA and HchA, and an example of the chromatogram from the enzyme assays.

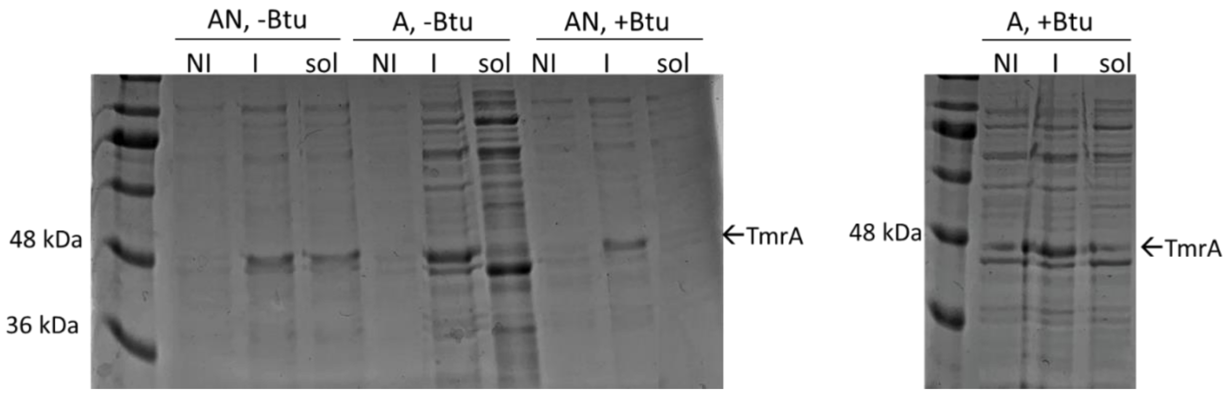

**Figure S1. SDS-PAGE of TmrA expression condition trials**. TmrA (48 kDa) expressed in E. coli under aerobic (A) or anaerobic (AN) conditions, and with (+ Btu) or without (- Btu) expression of pBAD42-BtuCEDFB. NI = not-induced (pre-induction), I = induced (whole cell), Sol = soluble lysate fraction. Ladder is Frogga Bio BLUeye pre-stained protein ladder.

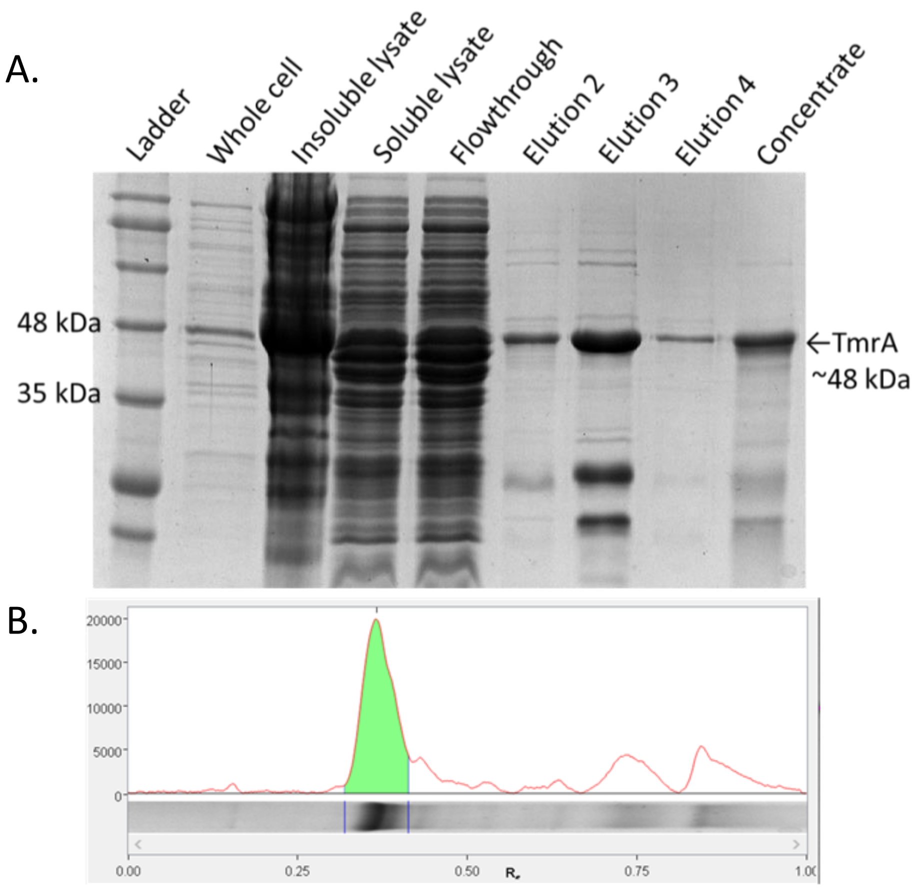

| Band No. | Band Label | Mol. Wt. (KDa) | Relative Front | Adj. Volume (Int) | Volume (Int) | Abs. Quant. | Rel. Quant. | Band % | Lane % |
|---|---|---|---|---|---|---|---|---|---|
| 1 | | N/A | 0.367 | 89,545,248 | 283,170,420 | N/A | N/A | 100.0 | 50.1 |

| | |
|---|---|
| Band Detection | Automatically detected bands with custom sensitivity: 91 |
| Lane Background | Lane background subtracted with disk size: 7.8 |
| Lane Width | 7.98 mm |

**Figure S2. (A) SDS-PAGE of TmrA** (48 kDa) expression and purification by nickel affinity chromatography. The whole cell lane was taken 18 hrs post-induction, the cell pellet was lysed using BugBuster extraction reagent (Millipore) and the soluble and insoluble fractions were imaged. The protein was purified by gravity chromatography using 2 mL of Ni-NTA resin from Qiagen, the flowthrough was collected for imaging. The elution fractions were combined and concentrated by a 30 kDa cut-off Millipore filter tube (Concentrate). Ladder is Frogga Bio BLUeye pre-stained protein ladder. (B) Screenshot of the lane density calculation of the concentrated TmrA used to determine approximate purity of TmrA, image from Image Lab v6.1.

**Table S6 Protein concentrations of purified TmrA and HchA.**

| Enzyme | Crude protein amount (mg) | Purified protein amount (mg) | Crude protein conc. (mg/mL) | Purified protein conc. (mg/mL) | Protein yield (%) | Purity (%) |
|---|---|---|---|---|---|---|
| TmrA | 292 | 2.3 ± 0.2 | 9.75 | 7.5 ± 0.7 | 0.8 ± 0.1 | 50 |
| HchA | 234 | 2.7 ± 0.4 | 7.8 | 9 ± 1 | 1.0 ± 0.1 | 39 |

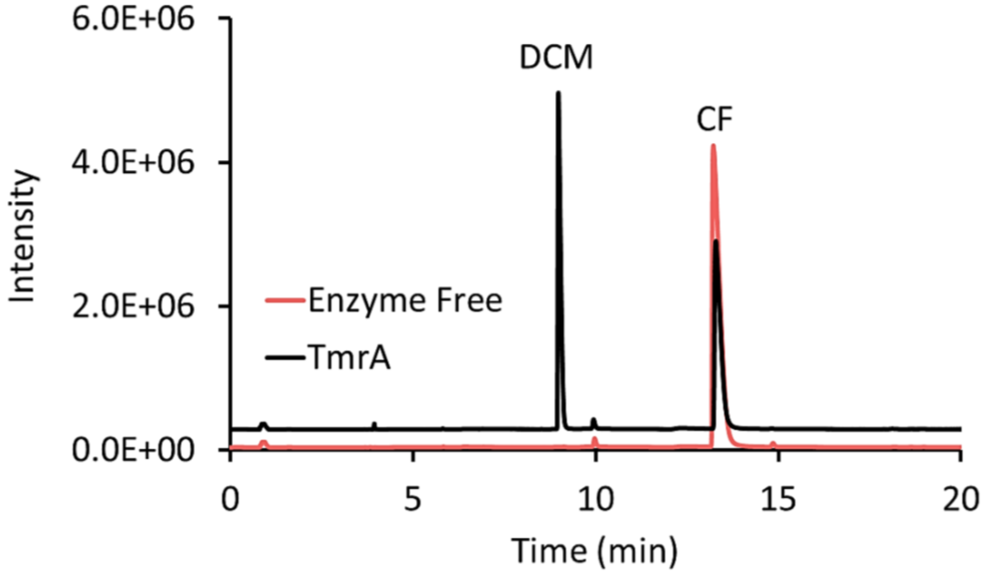

**Figure S3. Sample chromatogram** from GC-FID analysis of chloroform (CF) reduction by 3 µg TmrA into dichloromethane (DCM) after 1 hr of incubation at RT compared to the enzyme free negative control.

**Table S7. Raw data for TmrA lysates in Figure 2. (Excel)**

Table included in the accompanying excel document. The raw data for each replicate used in figures or references in the main text. CF = chloroform; DCM = dichloromethane; A = aerobic induction; AN = anaerobic induction; + Btu = with pBAD42-BtuCEDFB; - Btu = without pBAD42-BtuCEDFB. If strain is not specified E. coli ΔiscR is assumed. All assays were held for 24 hr under anaerobic conditions.

**Table S8. Raw data for TmrA, CfrA, and DcrA lysates in Table 1. (Excel)**

Table included in the accompanying excel document. Raw data for each replicate shown in Table 1 in the main text, or data relevant to Table 1. CF = chloroform; DCM = dichloromethane; 1,1,1-TCA = 1,1,1-trichloroethane; 1,1-DCA = 1,1-dichloroethane; 1,1-DCE = 1,1-dichloroethene; 1,1,2-TCA = 1,1,2-trichloroethane; 1,2-DCA = 1,2-dichloroethane; VC = vinyl chloride; CA = chloroethane. All assays were held for 24 hr under anaerobic conditions.

**Table S9. Raw data for HchA lysates in Table 2. (Excel)**

Table included in the accompanying excel document. The raw data for HchA for each replicate used in Table 2 in the main text, or data relevant to Table 2. Neither HCH substrate can be detected by gas chromatography. CF = chloroform; DCM = dichloromethane; 1,1,2-TCA = 1,1,2-trichloroethane; 1,2-DCA = 1,2-dichloroethane; VC = vinyl chloride; 1,1-DCA = 1,1-dichloroethane; CA = chloroethane; HCH = hexachlorocyclohexane; MCB = monochlorobenzene. All assays were held for 24 hr under anaerobic conditions.

**Table S10. Raw data for DHB14 and DHB15 lysates in Figure 3. (Excel)**

Table included in the accompanying excel document. The raw data for each replicate used in Figure 3B in the main text, or data relevant to Figure 3B. PCE = perchloroethene; TCE = trichloroethene; cDCE = cis-dichloroethene. All assays were held for 24 hr under anaerobic conditions.

**Table S11. Raw data for purified TmrA and HchA in Figure 4 and Figure 5. (Excel)**

Table included in the accompanying excel document. The raw data for each replicate used in Figure 4 and Figure 5 in the main text, or relevant data. . CF = chloroform; DCM = dichloromethane; 1,1,2-TCA = 1,1,2-trichloroethane; 1,2-DCA = 1,2-dichloroethane; VC = vinyl chloride; 1,1-DCA = 1,1-dichloroethane; CA = chloroethane; HCH = hexachlorocyclohexane; MCB = monochlorobenzene. All assays were held for 1 hr under anaerobic conditions. nkat is defined as nmol product/s.

#### SI Text 4 – *Dehalococcoides* sp. RDase Expression Attempts

The genes for VcrA (WP_012882535), BvcA (AAT64888), and TceA (AAW39060) were ordered in *pET-21* expression vectors from Twist Bioscience (San Francisco, CA, USA). The constructs were designed such that the RDase was truncated at the TAT cleavage site predicted by SignalP 5.0 and had a C-terminal x6His tag (52). The genes for VcrA and BvcA with their TAT signal peptide sequence had been previously amplified from genomic DNA isolated from the KB-1 mixed culture and cloned into *p15TV-L* with their TAT signal sequences by the Yakunin and Savchenko Labs (University of Toronto, Toronto, ON, Canada). The SUMO fusion enzymes were constructed by Gibson assembly of the PCR-amplified genes with extensions to introduce a 15 bp overlap sequence that were complimentary to the ends of the PCR-amplified *pET-SUMO* backbone, primers are found in Table S3. Descriptions of the expression vectors are found in Table S4. All plasmids were confirmed by Sanger sequencing using universal T7 primers and transformed into *E. coli* Δ*iscR* with *pBAD42-BtuCEDFB*.

The enzymes were expressed in *E. coli* Δ*iscR* with *pBAD42-BtuCEDFB* using the same protocol for small-scale expression and tested for activity as described in Materials and Methods. The culture lysates expressing the *pET-21* and *p15TV-L* constructs were tested for activity on trichloroethene (TCE), *cis*-dichloroethene (cDCE), and 1,2-dichloroethane (1,2-DCA). The mixed culture KB-1 grown on TCE as an electron donor was used as a positive control in all activity tests. No activity was seen from any of the heterologously expressed enzymes, but there was production of transformation products in the positive control. Multiple growth conditions were changed in an attempt to facilitate protein folding and solubility including the type of media, strain of *E. coli*, and the detergent in the lysis buffer. The enzymes were also co-expressed with chaperone proteins GroES/EL, trigger factor, and DnaK/DnaJ/GrpE, using commercial chaperone plasmid set from

Takara Bio Inc. (Shiga, Japan). Finally, we attempted expressing the enzymes as a fusion protein with an N-terminal SUMO solubility tag, which did increase the level of observed expression. The fusion proteins were assayed against the universal substrate 1,2-DCA only. None of these alterations resulted in detectable activity from the expression cultures indicating that the enzymes were not produced in their functional form.
